# Disease-causing mutations in subunits of OXPHOS complex I affect their physical interactions

**DOI:** 10.1101/526020

**Authors:** Gilad Barshad, Nicol Zlotinkov-Poznianski, Lihi Gal, Maya Schuldiner, Dan Mishmar

## Abstract

Mitochondrial complex I (C1) is the largest multi-subunit oxidative phosphorylation (OXPHOS) protein complex. Recent availability of a high-resolution human C1 structure, and from two non-human mammals, enabled predicting the impact of mutations on interactions involving each of the 44 C1 subunits. However, experimentally assessing the impact of the predicted interactions requires an easy and high-throughput method. Here, we created such a platform by cloning all 37 nuclear DNA (nDNA) and 7 mitochondrial DNA (mtDNA)-encoded human C1 subunits into yeast expression vectors to serve as both ‘prey’ and ‘bait’ in the split murine dihydrofolate reductase (mDHFR) protein complementation assay (PCA). We first demonstrated the capacity of this approach and then used it to examine reported pathological OXPHOS C1 mutations that occur at subunit interaction interfaces. Our results indicate that a pathological frame-shift mutation in the *MT-ND2* gene, causing the replacement of 126 C-terminal residues by a stretch of only 30 amino acids, resulted in loss of specificity in ND2-based interactions involving these residues. Hence, the split mDHFR PCA is a powerful assay for assessing the impact of disease-causing mutations on pairwise protein-protein interactions in the context of a large protein complex, thus revealing the mechanism underlying any associated pathogenicity.

## Introduction

The mitochondrial oxidative phosphorylation (OXPHOS) system is the main ATP production machinery in eukaryotic cells. The OXPHOS system comprises five multi-subunit protein complexes, of which NADH-ubiquinone oxidoreductase (complex 1, C1) is a major electron entry point into the electron transport chain (ETC) that is central to mitochondrial ATP synthesis. C1 dysfunction leads to devastating mitochondrial diseases ^1^ and is a hallmark of mitochondrial dysfunction in Parkinson’s disease ^2,3^, dementia ^4^ and the renal cancer oncocytoma ^5,6^. Moreover, C1 subunits are the most frequent gene targets for mitochondrial disease-causing mutations (reviewed by Koopman, et al. ^7^), some of which affect C1 assembly and stability in cultured cells and in patients ^8^. Hence, deciphering the molecular basis of C1 dysfunction is of major interest.

Mammalian C1 is the largest of the OXPHOS complexes (^∼^ 1 MDa), comprising 44 subunits ^9,10^. These include 14 core subunits conserved from bacteria to man ^11^, and 30 ‘super-numerary’ (accessory) subunits that were recruited during the course of evolution after the radiation of eukaryotes. Recently, high-resolution structural models of complex I from ovine ^12^ and bovine ^13^ hearts became available, as well as from human embryonic kidney cell line (HEK293) ^14^. Additionally, a deep view into the assembly process and assembly intermediates of C1 were recently reported ^15^. These studies enabled better understanding of the network of C1 subunit interactions. Nevertheless, most studies and experimental approaches taken to study C1 subunit interactions focused either on the holoenzyme or on assembly intermediates ^8,9,16-20^. As such, these studies allowed for assessing the impact of C1 pathological mutations only on the assembly process or on the overall function of C1 in certain cells ^21^, yet overlooked the possible impacts of mutations on specific subunit interactions. For this, a high-throughput screening method for considering all C1 pairwise interactions, is needed.

To create such an experimental platform, we cloned the entire set of human C1 protein subunits (N=44) into the two complementary fragments of murine dihydrofolate reductase (mDHFR) and performed a split-mDHFR protein complementation assay (PCA) in the yeast *Saccharomyces cerevisiae* ^22,23^. This experimental system enabled the detection of pairwise interactions between C1 subunits in a rigorous and experimentally facile manner. This strategy, moreover, provided a robust method to assess the impact of mutations on such interactions. To demonstrate the power of this approach, we first identified subunit interactions supported by available mammalian C1 structural models ^12-14^ and screened for pathological mutations that localize to subunit interaction interfaces. We then chose three changes, namely two SNPs and one frame shift mutation, all associated with mitochondrial dysfunction, and experimentally tested their effects. Our results indicate that a frameshift mutation in the mtDNA-encoded ND2 subunit, which underlies C1 dysfunction in a patient with severe exercise intolerance ^24^, led to a loss of specificity of ND2 subunit interactions. Our results provide proof-of-concept supporting the use of our experimental system as a resource to assess the impact of mutations on diverse protein-protein interactions in human C1.

## Results

### The split mDHFR protein complementation assay offers a robust platform for following protein-protein interactions in human complex I

To assess pairwise interactions between C1 subunits, we fused the coding sequences (CDSs) of all nuclear DNA (nDNA)- and mitochondrial DNA (mtDNA)-encoded human complex I subunits (N=44) to the 5’ end of that part of mDHFR gene encoding a C-terminal fragment (termed the C1-F3 constructs). In addition, 37 C1 subunits were fused to the 5’ end of that part of mDHFR gene encoding a N-terminal fragment (termed the C1-F1,2 constructs) (Fig. 1). In our hands, 8 subunits were not successfully cloned into the latter constructs (i.e, ND6, NDUFA9, NDUFA12, NDUFB9, NDUFS1, NDUFS4). While plasmids containing the C1-F3 constructs were inserted into MATa yeast, those plasmids containing the C1-F1,2 constructs were transformed into MATα yeast strains (Fig. 1, Supplementary Table S1) to enable rapid mating and the creation of diploids. The coding sequences of the mtDNA-encoded subunits (i.e., ND1-ND6 and ND4L) were edited to comply with yeast cytosolic codon usage. To reduce false-positive interactions, we excluded two highly over-expressed subunits (i.e., NDUFA2 and NDUFA5) from further analyses. To assess interactions between the various subunits, the haploid yeast strains were mated in a high-throughput manner (see Materials and Methods), thus creating 1,628 independent C1-F1,2 / C1-F3 diploid strains. The resultant colony sizes of these diploid strains were measured as a proxy for growth rate in the presence of methotrexate (MTX), thus selecting for yeast growth only when complementation of mDHFR activity had occurred (Fig. 1). This experiment revealed 94 significantly positive (feasible) pairwise interactions (Fig. 2). In seeking candidate subunit interactions able to report on the impact of pathological mutations, we focused on 19 interactions that were consistent with the C1 structural models from bovine and ovine heart ^12,13^ (Fig. 1). Such interactions were also consistent with recently published human C1 structures ^14^.

**Figure 1.**
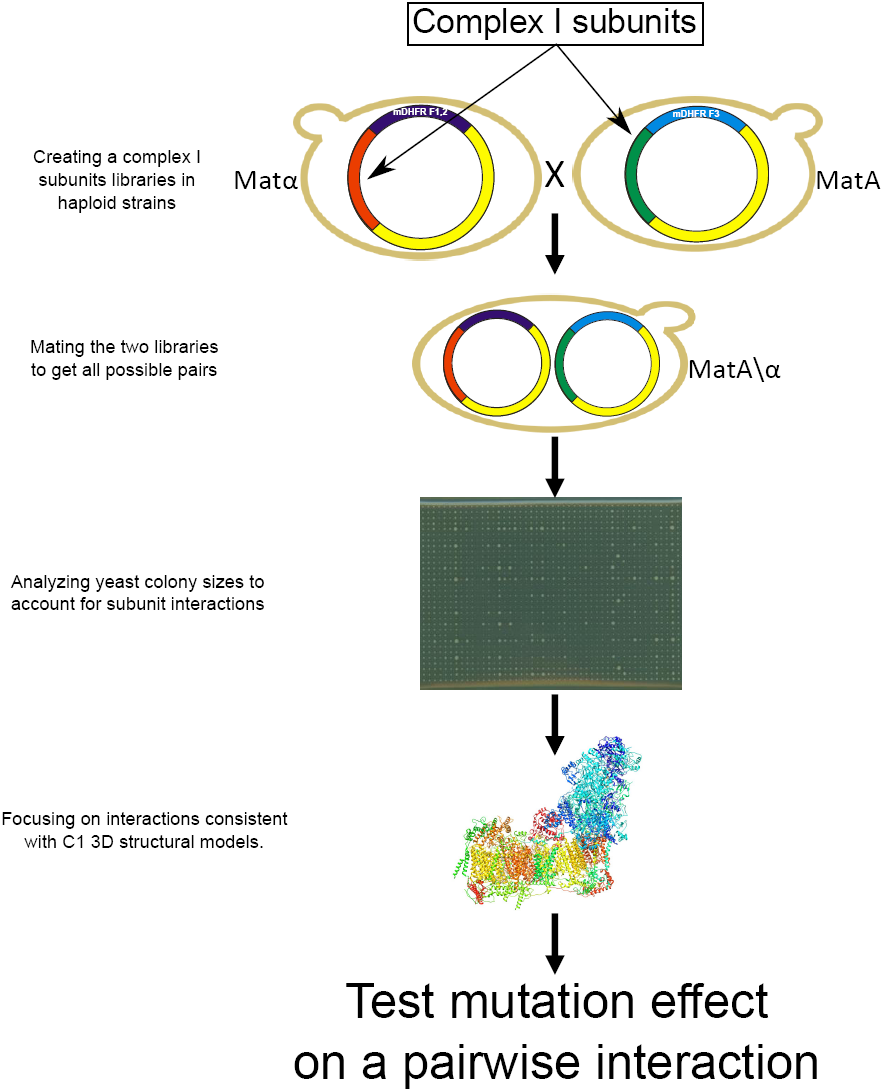
Workflow of the split-mDHFR-based screen for testing pairwise interactions between complex I subunits. Upper panel shows the construction of the haploid yeast split mDHFR libraries which harbor complex I subunit sequences, followed by mating, seeding on MTX+ plates and comparison to the structural data to identify testable interactions.

**Figure 2.**
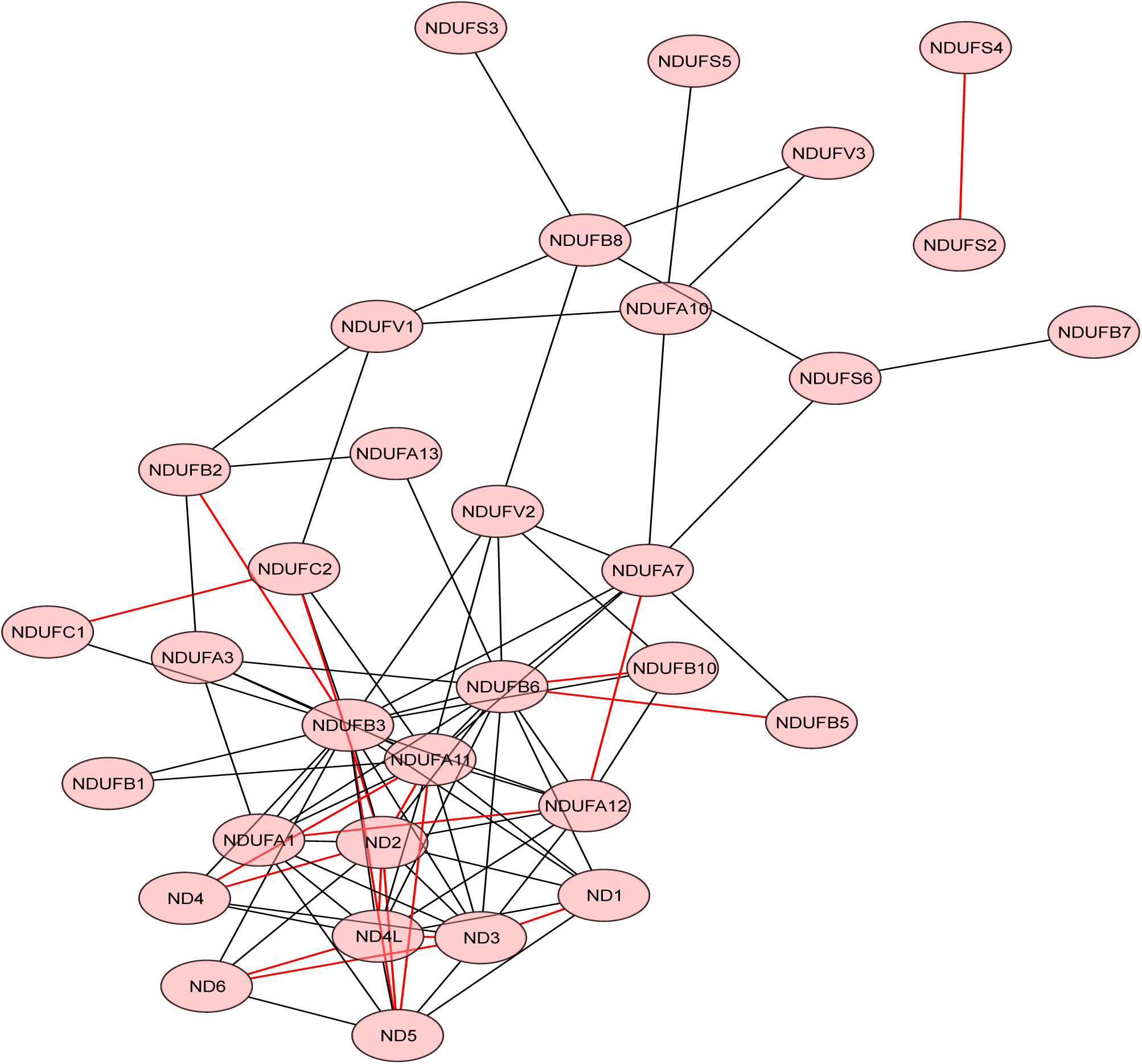
Network of identified C1 subunit interactions using the split mDHFR PCA. Edge length is positively correlated with the p-value for an interaction by a Mann Whitney U (MWU) test (see Materials and Methods). Red edges represent physical interactions that are consistent with the available 3D structural models of mammalian C1.

### Two disease-associated point mutations, affecting residues at the interaction interface of ND2 and ND5, had no effect on the pairwise subunit interaction

Next, we employed our assay to assess the impact of disease-associated human gene variants on physical interactions between the encoded proteins. First, we screened the literature for C1 disease-associated mutations that affect residues found at physical interaction interfaces of two C1 subunits, according to the available structures (Table 1, Supplementary File S1) ^25,26^. Two missense mutations at nucleotide positions 4917G and 4640A are associated with Leber’s hereditary optic neuropathy (LHON) in separate families ^25,27^. Both of the resulting amino acid substitution (i.e. N150D and I57M, respectively) occur in ND2 trans-membrane alpha-helixes (TMHs), which in the structure are adjacent to a ND5 subunit TMH, and hence potentially alter the interaction interface (Fig. 3A). Furthermore, the 4460A mutation was found in a Russian family from Novosibirsk who exhibited poor oxygen utilization in cybrids ^27^. The 4917G nucleotide substitution, an ancient mutation that defines the mtDNA genetic background haplogroup T, was associated with LHON only in the presence of the R2-JT haplogroup cluster-defining 4216C mutation ^25^. We introduced these two SNPs either separately or together into the ND2 re-coded sequence within the F3 mDHFR fragment-containing construct. We then mated the MATa yeast strains harboring these mutant constructs with a MATα strain harboring the ND5-F1,2 construct and conducted a dilution drop-test to assess the impact of these mutations on the ND2-ND5 interaction. Neither of the two single mutants showed any significant change in terms of yeast growth on selective media (containing MTX), suggesting that no significant impact on the ND2-ND5 interaction had occurred (Fig. 3B). Notably, the double mutant construct was not expressed to a detectable level in yeast, thus precluding any analysis of this combination of mutations (Fig. 3C, Supplementary Fig. S1).

**Table 1.**
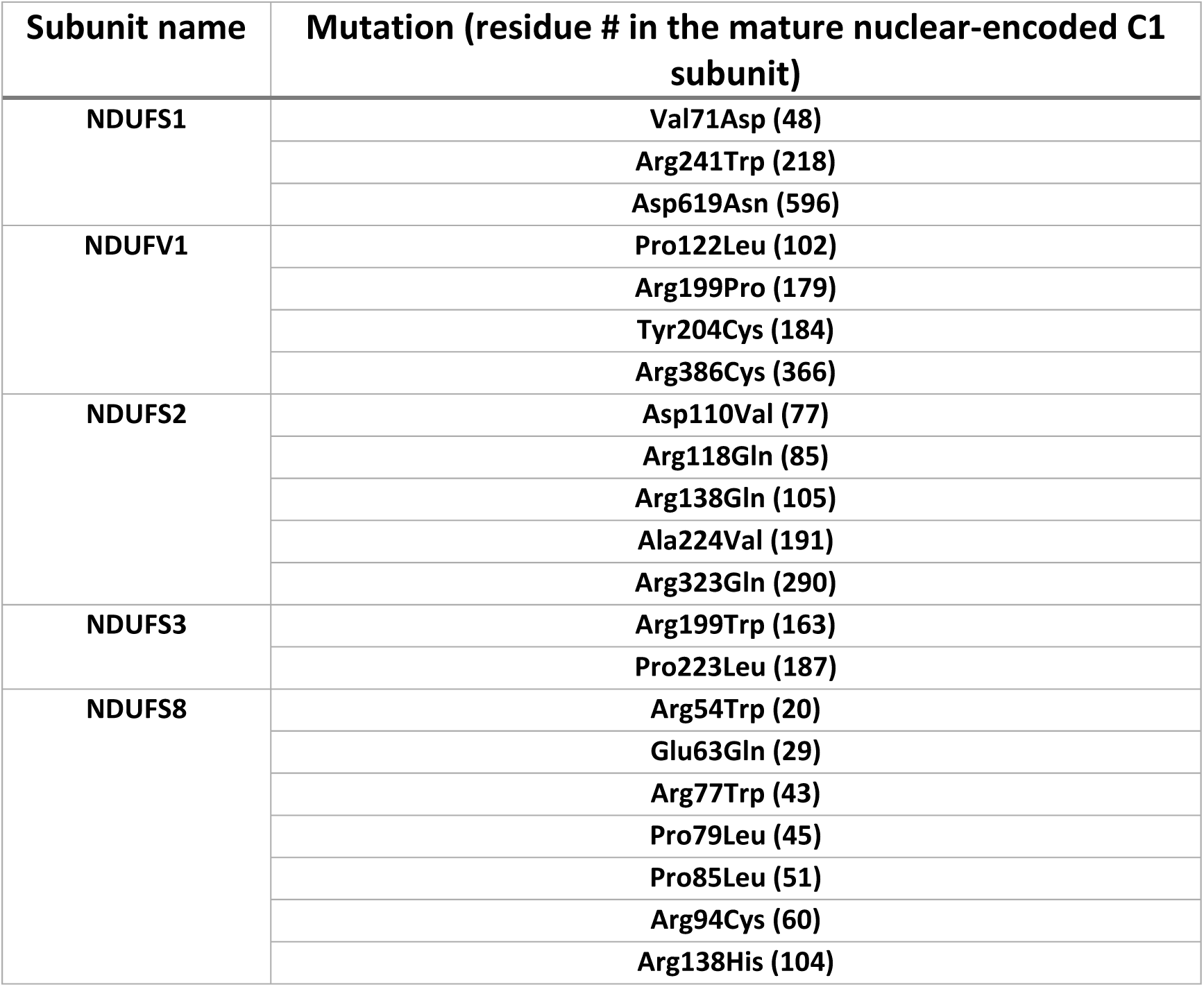
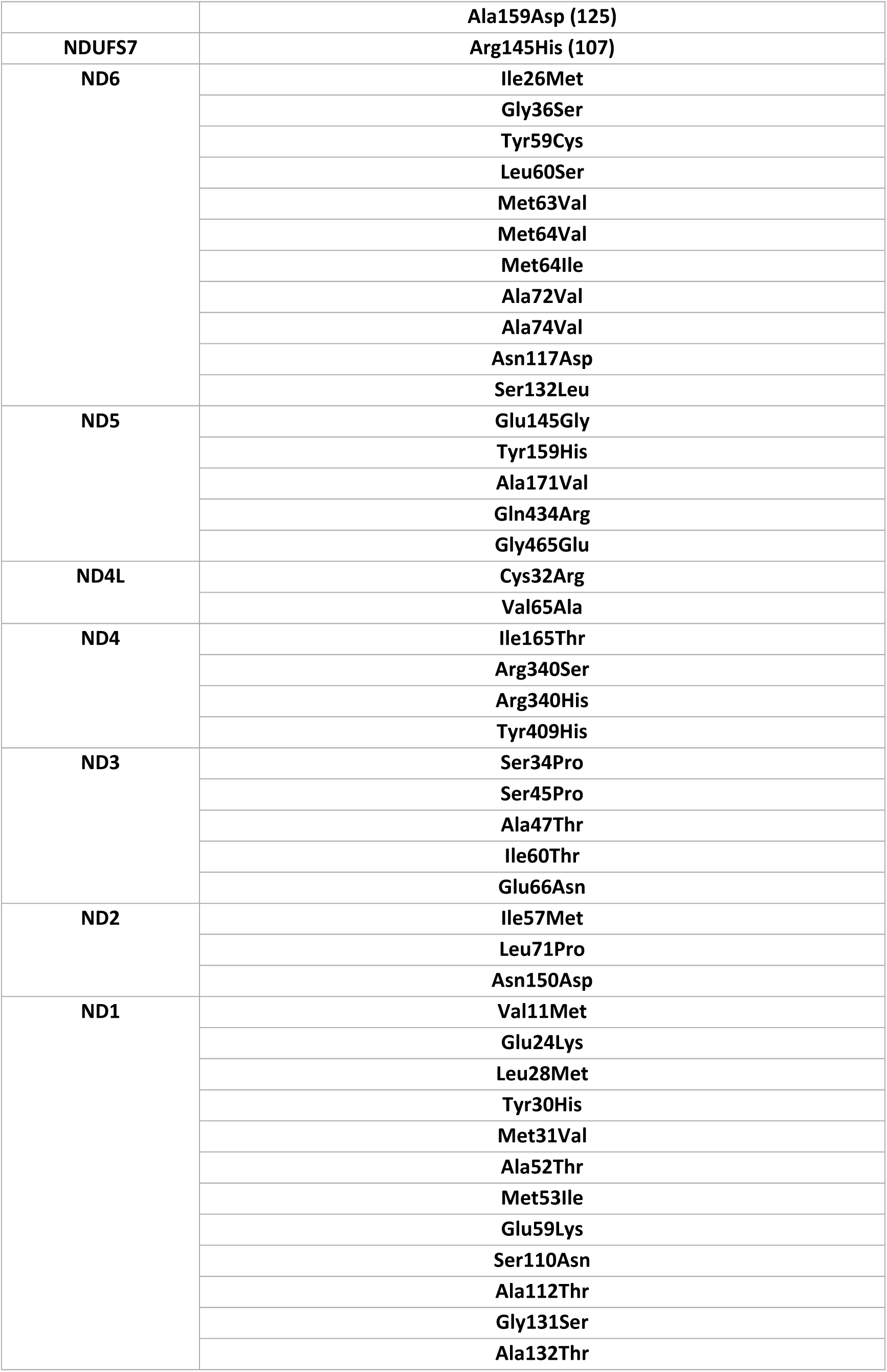
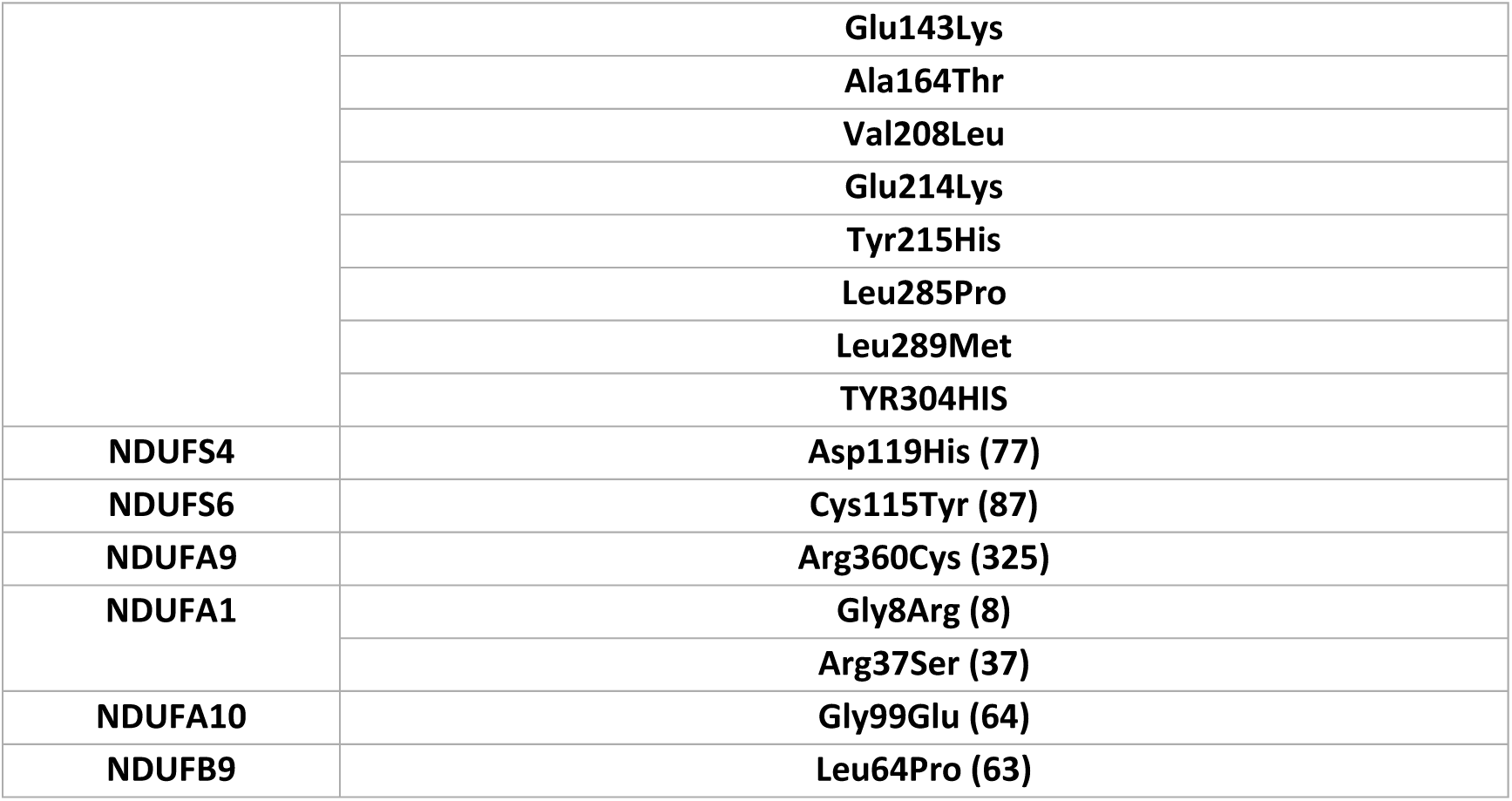
Single amino acid substitutions associated with C1 deficiency and potentially affecting protein-protein interactions between C1 subunits. The listed mutations were mostly extracted from ^26^ in addition to a complementary literature search.

| Subunit name | Mutation (residue # in the mature nuclear-encoded C1 subunit) |
| --- | --- |
| NDUFS1 | Val71Asp (48) |
|  | Arg241Trp (218) |
|  | Asp619Asn (596) |
| NDUFV1 | Pro122Leu (102) |
|  | Arg199Pro (179) |
|  | Tyr204Cys (184) |
|  | Arg386Cys (366) |
| NDUFS2 | Asp110Val (77) |
|  | Arg118Gln (85) |
|  | Arg138Gln (105) |
|  | Ala224Val (191) |
|  | Arg323Gln (290) |
| NDUFS3 | Arg199Trp (163) |
|  | Pro223Leu (187) |
| NDUFS8 | Arg54Trp (20) |
|  | Glu63Gln (29) |
|  | Arg77Trp (43) |
|  | Pro79Leu (45) |
|  | Pro85Leu (51) |
|  | Arg94Cys (60) |
|  | Arg138His (104) |
|  | Ala159Asp (125) |
| NDUFS7 | Arg145His (107) |
| ND6 | Ile26Met |
|  | Gly36Ser |
|  | Tyr59Cys |
|  | Leu60Ser |
|  | Met63Val |
|  | Met64Val |
|  | Met64Ile |
|  | Ala72Val |
|  | Ala74Val |
|  | Asn117Asp |
|  | Ser132Leu |
| ND5 | Glu145Gly |
|  | Tyr159His |
|  | Ala171Val |
|  | Gln434Arg |
|  | Gly465Glu |
| ND4L | Cys32Arg |
|  | Val65Ala |
| ND4 | Ile165Thr |
|  | Arg340Ser |
|  | Arg340His |
|  | Tyr409His |
| ND3 | Ser34Pro |
|  | Ser45Pro |
|  | Ala47Thr |
|  | Ile60Thr |
|  | Glu66Asn |
| ND2 | Ile57Met |
|  | Leu71Pro |
|  | Asn150Asp |
| ND1 | Val11Met |
|  | Glu24Lys |
|  | Leu28Met |
|  | Tyr30His |
|  | Met31Val |
|  | Ala52Thr |
|  | Met53Ile |
|  | Glu59Lys |
|  | Ser110Asn |
|  | Ala112Thr |
|  | Gly131Ser |
|  | Ala132Thr |
|  | Glu143Lys |
|  | Ala164Thr |
|  | Val208Leu |
|  | Glu214Lys |
|  | Tyr215His |
|  | Leu285Pro |
|  | Leu289Met |
|  | TYR304HIS |
| NDUFS4 | Asp119His (77) |
| NDUFS6 | Cys115Tyr (87) |
| NDUFA9 | Arg360Cys (325) |
| NDUFA1 | Gly8Arg (8) |
|  | Arg37Ser (37) |
| NDUFA10 | Gly99Glu (64) |
| NDUFB9 | Leu64Pro (63) |

**Figure 3.**
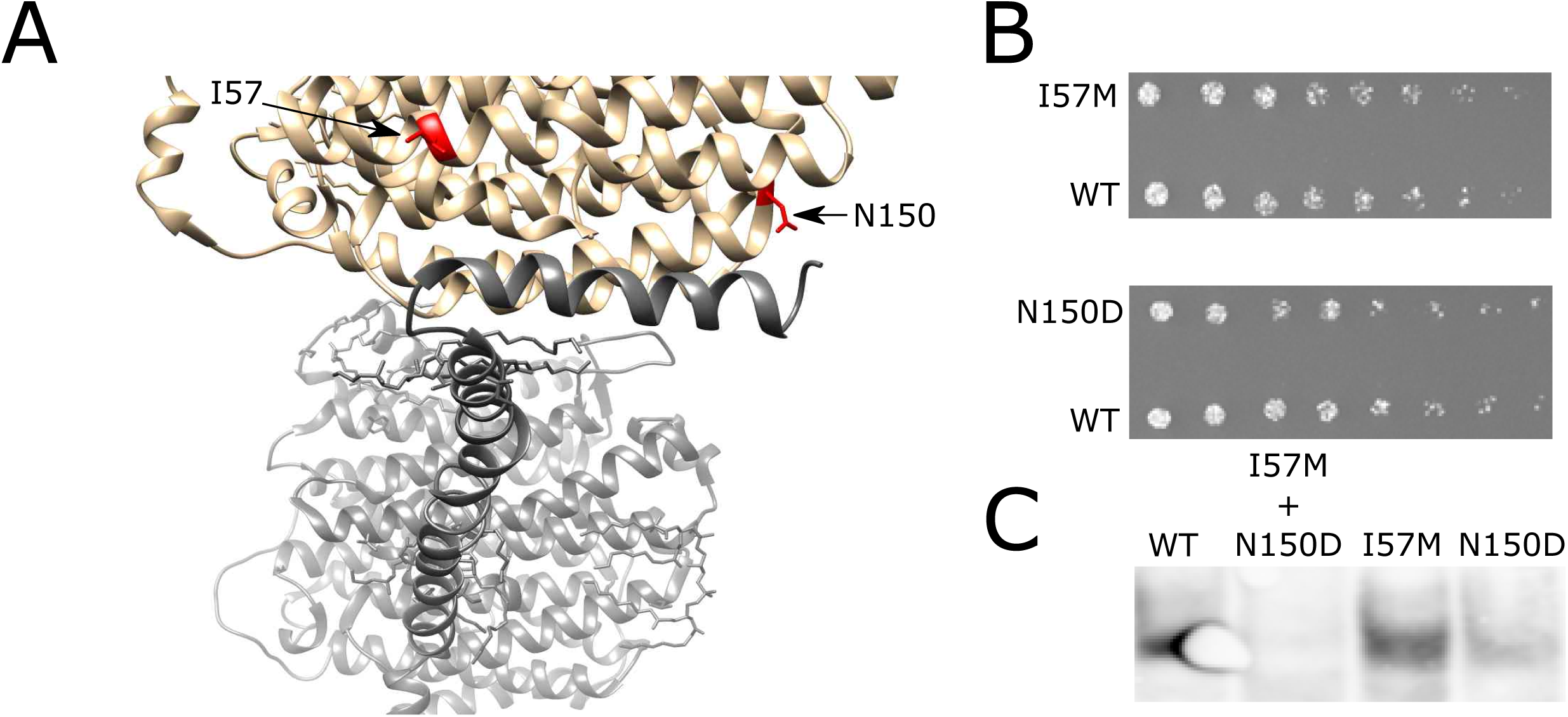
Ancient mtDNA SNPs which associate with LHON show no effect on the ND2-ND5 interaction. (A) The location of changes in the ND2 sequence (red) in two TMHs (gray) and at the interaction interface with ND5 (black). (B) Dilution drop-test results for the two ND2 mutants against wild type (WT) ND2. (C) Western blot demonstrating expression of the two single mutants and WT ND2 but not of the double mutant.

### A two base-pair deletion causing an *MT-ND2* frame shift mutation leads to the emergence of interaction promiscuity

We next examined the impact of a double adenine deletion within an adenine triplet at mtDNA nucleotide positions 5132 to 5134, causing a frame shift in the *MT-ND2* sequence. This deletion was reported in a 28-year-old patient with mitochondrial myopathy and severe exercise intolerance ^28^. Such frameshift led to replacement of the 126 C-terminal residues harboring three TMHs in ND2 by 30 highly hydrophilic residues (Fig 4A). We cloned the ND2 deletion mutant into the ND2-F3 construct and introduced the resulting plasmid into the MATa yeast strain, which was in turn mated with MATα strains containing one of four ND2 interactors (i.e., NDUFC2, ND4, ND4L and ND5) (Fig. 4B) fused to the F1,2 mDHFR fragments. Here too, we performed dilution drop-test experiments for all four mating-pairs and compared the relative growth rates of the diploid strains containing the ND2 mutants with that of diploid strains harboring wild type ND2 (Fig. 5A). Our results indicated unexpected increased growth rates of the ND2 mutant strains when mated with NDUFC2- and ND4-expressing strains, while no significant change in growth rate was evident when the ND2 mutant strain was mated with ND4L or ND5 (Fig. 5B). However, while inspecting the control diploid strains, harboring mutated *ND2* and an empty (i.e., not encoding any C1 subunits) F1,2 construct, we noticed much higher growth as compared to the diploid strain harboring wild type ND2 and the empty F1,2 vector (Fig. 5B). This suggests that the ND2 mutant had become a promiscuous interacting protein.

**Figure 4.**
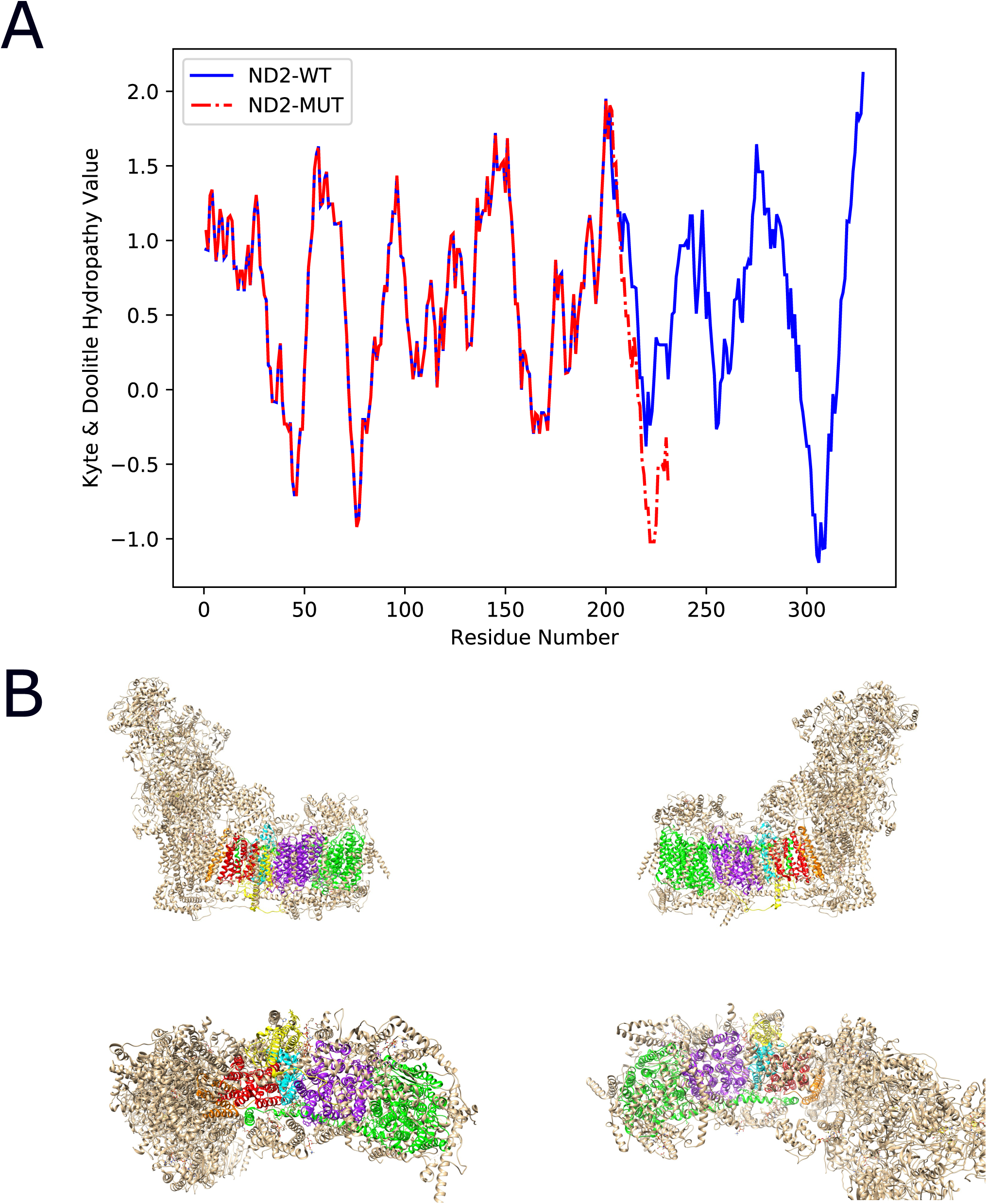
A disease-causing ND2 frameshift mutant results in a shorter, more hydrophilic C-terminus. (A) Kyte-Doolittle hydropathy plot comparing the hydrophobicity of the wild type (blue) and the ND2 frameshift mutant (red). (B) Four viewpoints of human C1. Red - the N- terminal region of ND2, cyan - the deleted ND2 C-terminal region. The four subunits that interact with ND2: NDUFC2 (yellow), ND4 (purple), ND5 (green) and ND4L (orange).

**Figure 5.**
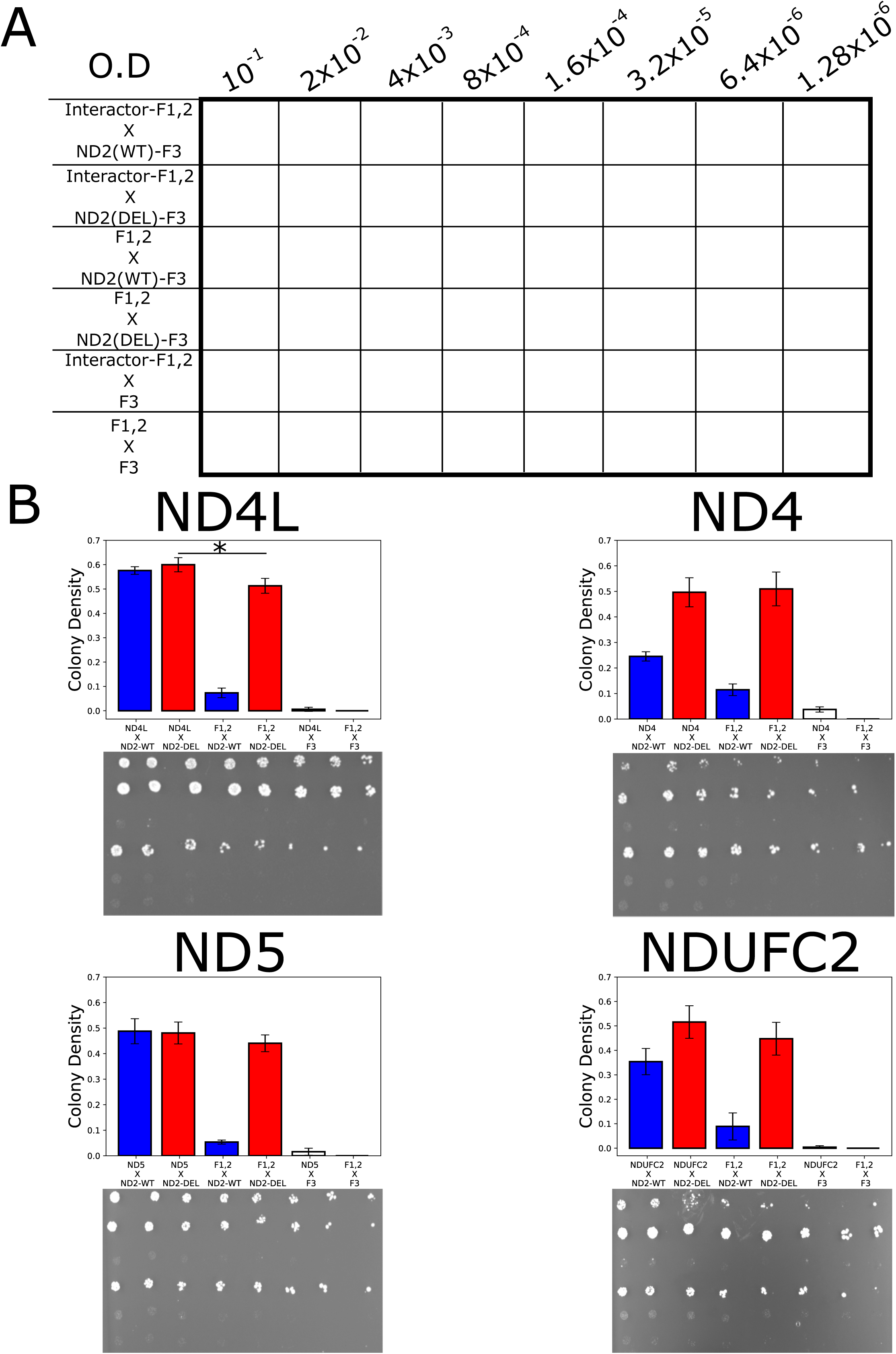
The ND2 frameshift mutation results in promiscuous C-terminal interactions. (A) The yeast growth plate designed for the dilution drop-test. ND2-DEL – frameshift mutant; ND-WT – wild type ND2. (B) Representative drop-test plates and histograms summarizing the yeast colony intensity obtained in three biological replicates.

However, promiscuity did not explain the increase in interaction strength in all cases; ND4L exhibited a stronger interaction with the mutant ND2, as compared to the empty vector control (Fig. 5B). A close inspection of the C1 structure revealed ND4L to be the only subunit tested that interacts with the N-terminal portion of ND2 (Fig. 4B). This suggests that the specific interaction of ND2 and ND4L, which is dictated by their interaction interface, is not affected by the deletion mutation in *ND2*.

## Discussion

The availability of structures of multi-subunit protein complexes enables predicting the potential impact of mutations on subunit interactions. Such predictions can serve to design experiments that test the impact of such mutations. Nevertheless, such experiments cannot easily employ classical structural biology methodologies, which are not compatible with high-throughput approaches and are less accessible to most molecular biologists. Hence, we established an assay to enable such assessment to be carried out easily. Specifically, we used the split mDHFR protein complementation assay (PCA) to assess the impact of mutations at interaction interfaces between C1 subunits on protein-protein interactions. First, we cloned all known human C1 subunits (N=44) into constructs harboring either of both parts of split mDHFR, thus establishing a resource which enables testing for all pairwise interactions. Such a resource offers the obvious advantage of accessibility and relative ease-of-use, relying on an assay based on yeast mating and measurement of growth rate on selective medium (containing methotrexate). The main disadvantage, especially in the case of a multi-subunit protein, is that the PCA is limited to assessment of pairwise interactions. Additionally, in OXPHOS C1, several subunits are simultaneously added during each stage of the assembly process ^29,30^. This suggests that many interactions between subunit pairs likely depend on the presence of additional subunits; naturally, an assay designed to test pairwise interactions is blind to such interactions. This may explain, at least in part, why only 19 interactions in our assay were consistent with the 3D structures. Regardless, our observed interactions contained core subunits, which play a major role in the activity of C1 and reflect the usefulness of our tool for assessing the impact of mutations on such interactions.

As proof-of-concept, we focused our experimental assessment on three changes, considering two disease-associated SNPs and one frameshift mutation, all occurring in the mtDNA-encoded ND2 subunit. The first SNP, 4640A, was originally found in a Russian family from Novosibirsk. The 4640A mutation resulted in poor oxygen utilization by cybrids, although these experiments did not reveal any reduction in the specific activity of C1, relative to control lymphoblast cells ^27^. Close inspection of this mutation showed that it altered an amino acid position with poor evolutionary conservation. Additionally, the 4640A mutation appears to associate with the U3b sub-haplogroup in the human mtDNA phylogenetic tree. Similarly, the 4917G mutation associates with both the T and N1b haplogroups. Haplogroup T associates with LHON ^31^, and with reduced sperm motility in the Spanish population ^32^ and haplogroup N1b altered the susceptibility of Ashkenazi Jewish type 2 diabetes patients to develop complications ^33^. Notably, haplogroup T cybrids possess a higher capability to cope with oxidative stress in the form of an H_2_O_2_ challenge ^34^. Taken together, the impact of SNPs that co-occur with certain mtDNA genetic backgrounds cannot be easily distinguished from the linked set of mutations. One should thus take into consideration the fact that our observed lack of impact of both the 4640 and 4917 mutations on C1 subunit interactions may either attest to their minor functional effect, suggesting that their impact is not manifested at the level of protein-protein interactions, or that this reflects the resolution limitations of the PCA. These possibilities cannot be resolved at this point and requires additional experiments.

Unlike the tested point mutations, a pathological *MT-ND2* frameshift mutation had a dramatic impact on the pattern of subunit interactions. Our results suggest that the *MT-ND2* frameshift mutant generated a promiscuous interacting protein, specifically in interactions involving the region encoded downstream to the frameshift. Such a molecular phenotype might explain the appearance of the disease at the relatively high (90%) heteroplasmy level seen in the patient affected by the deletion mutation ^24^. It is also possible that at high heteroplasmy levels, the translation product of the mutated *MT-ND2* gene interacts with many members of the mitochondrial proteome, thus interfering with the proper ND2 interaction pattern, thereby causing the mitochondrial dysfunction observed in the clinic. Additionally, our results suggest that interaction of the N-terminal region of ND2 with the ND4L subunit was not affected, suggesting that the promiscuous interaction phenotype is limited to interactions involving the C-terminal region of ND2, thus excluding the possibility of a general mis-folding effect of ND2. These results provide, for the first time, a molecular mechanistic explanation for the pathophysiology of the *ND2* frameshift mutation.

As C1 structure and assembly pathway are very complex, studies of the phenotypic impact of human disease-causing mutations and disease-associated polymorphisms have largely relied on previous measurements of C1 enzymatic activity ^2,27,35^ or on earlier work on the assembly of the holoenzyme. Still, little is known to date of the effects of human mutations on specific pairwise interactions between C1 subunits. We speculate that protein-protein interactions between C1 subunits that occur independently of additional subunits are important for the structure and proper assembly of C1.

A recent RNA-seq analysis from multiple human tissues revealed that certain C1 subunits show tissue-dependent expression patterns ^36^, thus questioning the generality of structural analysis of the complex from specific tissues, such as the heart or liver. With this in mind, our analysis of C1 subunit interactions is not linked to a particular tissue, and hence is less influenced by tissue-dependent subunit composition considerations.

In summary, we provide a resource of clones for all known human C1 subunits ready to be tested for pairwise protein-protein interactions using the split mDHFR PCA. Our experimental assessment of the impact of mutations on pairwise C1 subunit interactions serve as a proof-of-concept and may encourage using this approach to study the impact of specific mutations on other multi-subunit protein complexes, such as the ribosome. Our finding of a profound impact of a *MT-ND2* frameshift mutation on interactions with partners revealed in the crystal structure of the complex provides the first direct explanation of the molecular mechanism underlying the phenotypic impact of this mutation. This discovery increases the motivation to recruit the system established in the current study to screen for the impact of other mutations of interest on confirmed subunit interactions.

## Materials and methods

### Yeast growth media

The complete recipes for yeast media used in the current study are detailed in Supplementary Table S2.

### Preparation of split mDHFR libraries

Two 2-micron plasmids were used for human C1 subunit library preparation. Plasmid p340 includes amino acid sequence-coding motifs at the cloning site for a linker peptide (GlyGlyGlyGlySer)X2 followed by the ORF of the yellow fluorescent protein (YFP)-coding gene, followed by another (GlyGlyGlyGlySer)X2 peptide linker and DNA encoding mDHFR N’ terminal fragments (F1,2). Expression of this fusion construct was under the transcriptional control of the *GAP* promoter and *CYC1* terminator. Plasmid pMS141 includes amino acid sequence-coding motifs at the cloning site for a linker peptide (GlyGlyGlyGlySer)X2 followed by DNA encoding a mDHFR C’ terminal fragment (F3), and is expressed under the transcriptional control of the *TEF* promoter and *ADH1* terminator. In both plasmids, a *SpeI* restriction site was fused to the 5’ codon of the constructs (Supplementary Fig. S2).

ORFs of nDNA-encoded human C1 subunits were amplified from commercial human cDNA samples (See Supplementary Table S3 for the list of sources of human RNA). ORFs of the mtDNA–encoded subunits were recoded to match the cytoplasmic translation code (Supplementary Table S4), generated commercially according to our design and cloned into pUC57 plasmids (OriGene). To create the two libraries, namely a F3 library in strain BY4741 [MATa, *his3Δ1, leu2Δ0, met15Δ0, ura3Δ0*] and a YFP-F1,2 library in strain BY4742 [MATα, *his3Δ1, leu2Δ0, lys2Δ0, ura3Δ0*], the ORF encoding each subunit was amplified using primers harboring 5’ tails of ^∼^50 bp homology to either plasmid p340 or pMS141 (primers are listed in Supplementary Table S5). PCR amplification of the inserts was performed using Phusion DNA polymerase (Thermo-Scientific), with the following amplification protocol: Two min at 98°C followed by 35 cycles of 30 sec at 98°C, 30 sec at the melting temperature of the primers used and 72°C for an elongation time calculated as template length in kilo base pairs multiplied by 30 sec. To perform homologous recombination-based cloning in yeast, 15 µL of the PCR reaction were used for co-transformation with the relevant plasmid into yeast haploid strains (plasmids p340 and pMS141 were used for transformation into the BY4742 and BY4741 strains, respectively). To improve the efficiency of the cloning process in yeast, the plasmids were linearized by digestion with *SpeI*, and 200 ng of the digested plasmids were used for co-transformation with the PCR fragments. The yeast transformations were performed as previously described ^37^. Following co-transformation of the inserts and the linearized plasmids, the yeast were grown on selective plates, namely SD–His agar plates for plasmid p340-transformed cells and YPD agar plates with cloNAT for plasmid pMS141-transformed cells, for a minimum of 48 h at 30°C. Colonies (N=8 to 10) were picked and streaked onto a similar second selective plate and grown overnight at 30°C.

To validate successful cloning, a crude extract containing plasmid DNA was prepared by boiling small patches of the streaked colonies in 50 µL of 20 mM NaOH at 100°C for 25 minutes and centrifugation for 2 min at 2,500xg. The supernatant was used as template in a 10 µL PCR amplification using primers from the plasmid sequence flanking the insert (see Supplementary Table S6). PCR products of appropriate size were Sanger-sequenced at the Ben-Gurion University sequencing unit. The sequences obtained were compared with the RefSeq sequences of the subunit-encoding ORFs (see Supplementary Table S7 for C1 subunits RefSeq protein accession numbers).

### Split mDHFR screen

All 44 human C1 subunits were cloned into plasmid pMS141 (mDHFR fragment F3) and 37 of the subunits were also cloned into plasmid p340 (mDHFR fragments F1,2). To test for all possible pairwise interactions between C1 subunits, seven haploid strain selection plates were prepared, harboring 1,536 colonies each. Three SD-His agar plates were prepared for plasmid p340-transformed BY4742 cells and four YPD agar plates were prepared with cloNAT for plasmid pMS141-transformed BY4741 cells. The haploid strain libraries were seeded onto haploid selection plates and grown at 30°C for 48 h, replicated on a second identical haploid selection plate and grown for an additional 24 h to reduce contamination. The libraries were then mated so that each of the plasmid p340-transformed BY4742 clones (three 1,536 clone plates) could be crossed with each of the four plasmid pMS141-transformed BY4741 plates, and vice-versa, yielding a total of 12 mating plates.

Mating was performed while growing the yeast on non-selective YPD agar plates at 30°C overnight to allow for the differently selected haploid strains to mate. These plates were subsequently replicated onto diploid selection SD-His agar plates with cloNAT. The resulting yeast colonies were grown overnight at 30°C and then, to reduce haploid contamination, were replicated onto a second diploid selection plate and once again grown overnight at 30°C. Each of the 12 diploid selection plates from the second round of diploid replication were replicated onto SD agar plates containing MTX and placed in a 30°C incubator for 5 days. All yeast plate replicates and mating experiments were performed using a RoToR bench top colony arrayer (Singer Instruments, UK).

Plate distribution of the haploid strains was designed so that crosses of the BY4741 and BY4742 strains would result in a minimum of 9 replicates of each of tested subunit A-F1,2 x subunit B-F3 interactions, with an additional 15 colonies representing the control population for each tested interaction. Here, 9 subunit A-F1,2 x F3 colonies did not contain an insert and 6 colonies were F1,2 without insert x subunit B-F3 crosses. Each haploid plate harbored control colonies in which the construct did not include an insert.

### Statistical analysis of the yeast colonies

Colony area size was measured using Balony ^38^. Firstly, colony sizes were normalized to the median size of all colonies within their row and column on each plate. Then, a one-tailed Mann Whitney U (MWU) test was performed with the alternative hypothesis of the form P(case>control) > P(control>case). An interaction was considered positive only if the ‘case’ group was significantly larger than both the ‘control’ groups described above (i.e., colonies resulting from mating of subunit A-F1,2 x F3 without insert and from F1,2 without insert x subunit B-F3). When colony area size was equal to zero, the colony was excluded from further statistical analysis to avoid technical bias.

### Drop test for evaluating the effect of the *ND2* frameshift mutation

Haploid strains were seeded onto haploid selection plates (SD–His agar plates for strain BY4742 containing the F1,2 constructs with DNA encoding ND4, ND4L, ND5 or NDUFC2 or insert-less and YPD agar plates with cloNAT for strain BY4741 containing the F3 constructs with DNA encoding ND2-WT, ND2-DEL (frameshift mutant), ND2-I57M, ND2-N150D or ND2-I57M + N150D or insert-less), grown at 30°C for 48 h, replicated onto a second identical haploid selection plate and grown for an additional 24 h to reduce contamination. Mating was performed while growing the yeast on non-selective YPD agar plates at 30°C overnight. These plates were subsequently replicated on diploid selection SD-His agar plates with cloNAT. The resulting yeast colonies were grown overnight at 30°C and then, to reduce haploid contamination, were replicated onto a second diploid selection plate and once again grown overnight at 30°C. After validating expression of the ND2 variants, the diploid strains were grown in SD liquid medium containing MTX (to reduce nucleotide pools) and SD liquid medium containing DMSO (control) for 24 h at 30°C with shaking. These yeast strains were diluted to OD_600_ = 0.1 in fresh SD liquid medium containing MTX or DMSO, respectively, and each of the diluted strains was further diluted 7 times at a 1:5 ratio. Five µL of each diluted diploidic strain were placed on a SD agar plate containing MTX or DMSO and placed in a 30°C incubator for 5 days. Images were taken at days 2-5 to assess colony intensity and to avoid reaching a growth asymptote. The overall highest differences between colonies at the same dilution resulting from the different matings were spotted on day 3 with the smallest intra-mating variations at the seeding density of OD_600_ = 0.1. As the only measurable background was upon mating the insert-less-F1,2 control with either ND2-WT- or ND2-DEL-expressing cells, we tested for significant growth over background by comparing the growth of the interactor-F1,2 X ND2-WT/ND2-DEL cross with the appropriate control mating of the insert-less-F1,2 X ND2-WT/ND2-DEL cross, respectively. Such pairwise comparisons were made using a two-tailed, unpaired Student’s t-test with equal variance.

### Validation of F3 fragment expression by western blot

Diploid strain colonies representing all matings performed for the drop-test were picked from the second diploid selection plates (see above) and grown overnight in SD-His liquid medium supplemented with cloNAT at 30°C with shaking. The strains were diluted to OD_600_ = 0.1 and grown overnight in SD-His liquid medium supplemented with cloNAT until mid-log phase (OD_600_ = 0.6-0.8). Approximately 2.5 OD_600_ units (3.125-4.167 ml) of the cells were centrifuged at 1000g for 5 min, resuspended in 100 mM NaOH and incubated for 5 min at room temperature. The resulting cell extracts were centrifuged at 16,000xg for 3 min and the protein-containing pellet was resuspended in SDS-containing loading buffer and incubated at 70°C for 10 min. Thirty µg aliquots were loaded onto 4%-20% polyacrylamide gels and the separated proteins were transferred to a polyvinylidene difluoride membrane at ^∼^330 mA for 1 h. The membrane was then subjected to a standard blotting protocol with primary rabbit anti-mDHFR(F3) antibodies (Sigma-Aldrich D0942) diluted 1:1000 and horseradish peroxidase-conjugated goat anti-rabbit secondary antibodies. Antibody binding was visualized with an enhanced chemiluminescent solution (WesternBright Sirius, advansta, BGU) for 5 min and image analysis (LAS-3000 luminescent image analyzer, Fujifilm). Such analysis revealed an additional ^∼^ 10 kDa band in the ND2-WT strains that was absent from the ND2-DEL strains, probably reflecting some level of unfused F3 expression product (Supplementary Fig. S3).

## Acknowledgements

The authors wish to thank Prof. Raz Zarivach (BGU) for critical discussion during early stages of the manuscript. This study was funded by grants 372/17 and 610/12 from the Israeli Science Foundation, a Binational Science Foundation grant number 2013060 and a US army life sciences division grant (6793LS) awarded to DM. The Schuldiner lab is supported by a DIP grant (P17516). MS is an incumbent of the Dr. Gilbert Omenn and Martha Darling Professorial Chair in Molecular Genetics.

## Author Contribution

G.B. constructed the library of complex I subunits and performed the vast majority of the analyses (experimental and computational), and participated in writing the manuscript; N.Z-P. assisted in the split mDHFR assay and performed some of the western blots; L.G. assisted G.B. in performing the initial screens of protein-protein interactions; M.S. assisted in the experimental design and critical reading of the manuscript; D.M. conceived the idea and wrote the manuscript.

## Additional information - Competing Interests

The authors have no competing interests as defined by Nature Research, or other interests that might be perceived to influence the results and/or discussion reported in this paper.

## Supplementary Figure legends

**Figure S1. Western blot demonstrating expression of the two single mutants and WT ND2 but not of the double mutant - full size blot.** This figure corresponds to Figure 3C. Dashed rectangle shows the area from which Figure 3C was cropped.

**Figure S2. Maps of the constructs used in the split mDHFR PCA.** Human C1 subunit insertion sites in each plasmid are indicated.

**Figure S3. Validation of protein expression of the ND2 frameshift mutant and the wild type protein.** Western blot representing three independent mating experiments of either wild type ND2 or the ND2 frameshift mutant with all four tested interactors. Bands representing the wild type and the frameshift mutant are indicated. Complementary full western blot is shown at the lower panel. Dashed rectangle represent the area from which the upper panel was cropped.

