## Supplementary Figures and Tables for "Disease-causing mutations in subunits of OXPHOS complex I affect their physical interactions"

Supplementary Figure S1

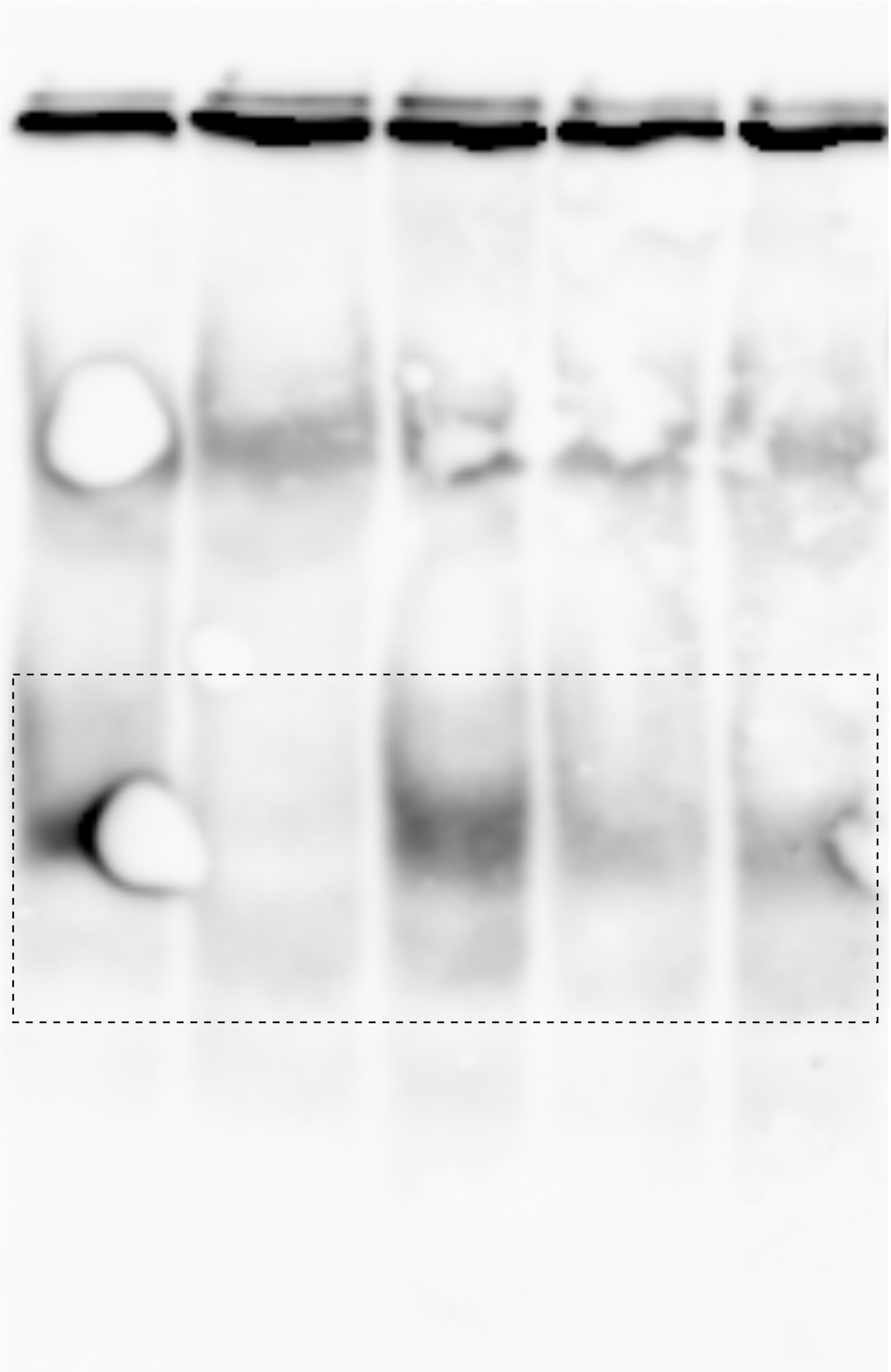

**Figure S1. Western blot demonstrating expression of the two single mutants and WT ND2 but not of the double mutant - full size blot.** This figure corresponds to Figure 3C. Dashed rectangle shows the area from which Figure 3C was cropped.

Supplementary Figure S2

A

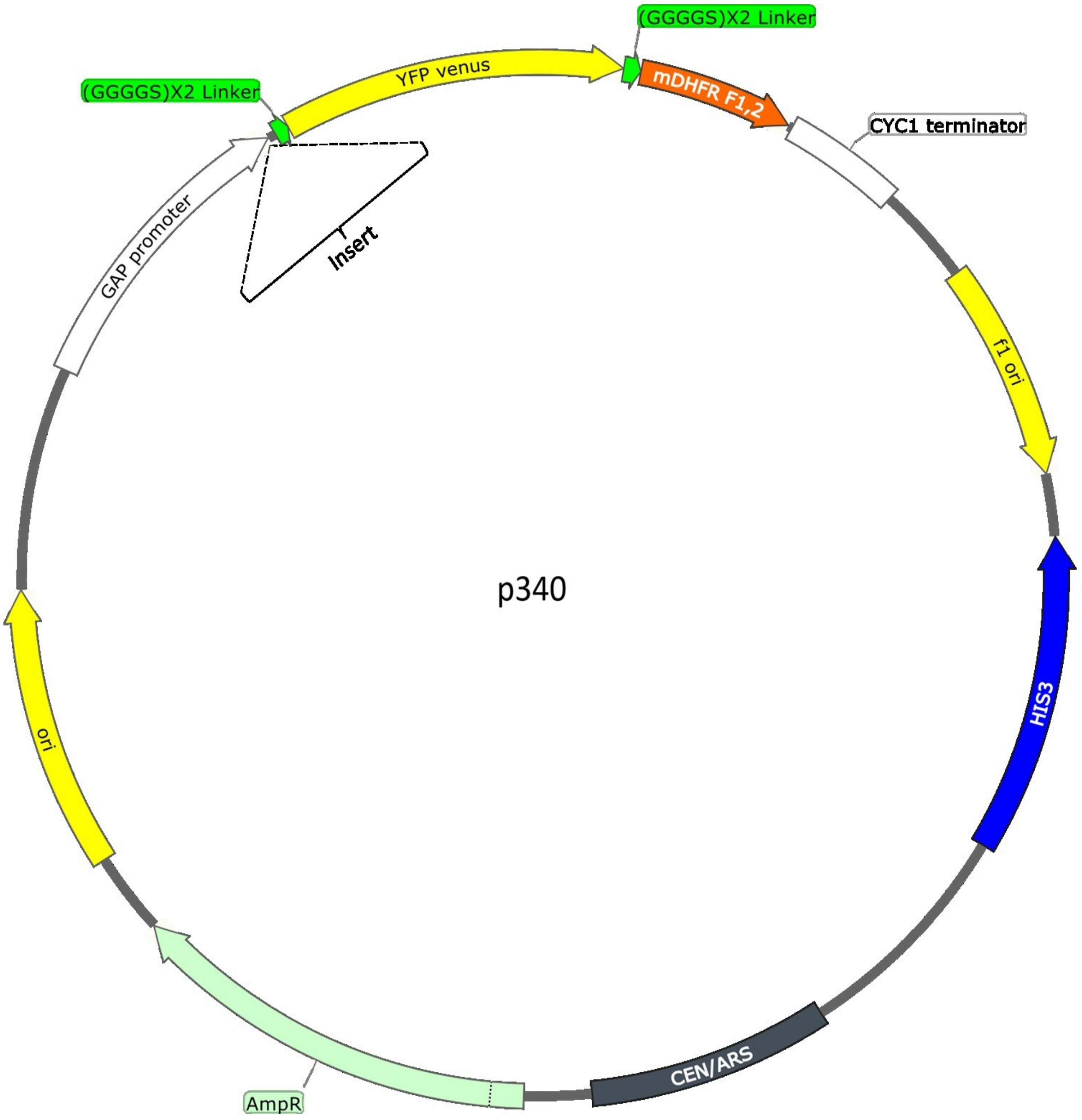

**B**

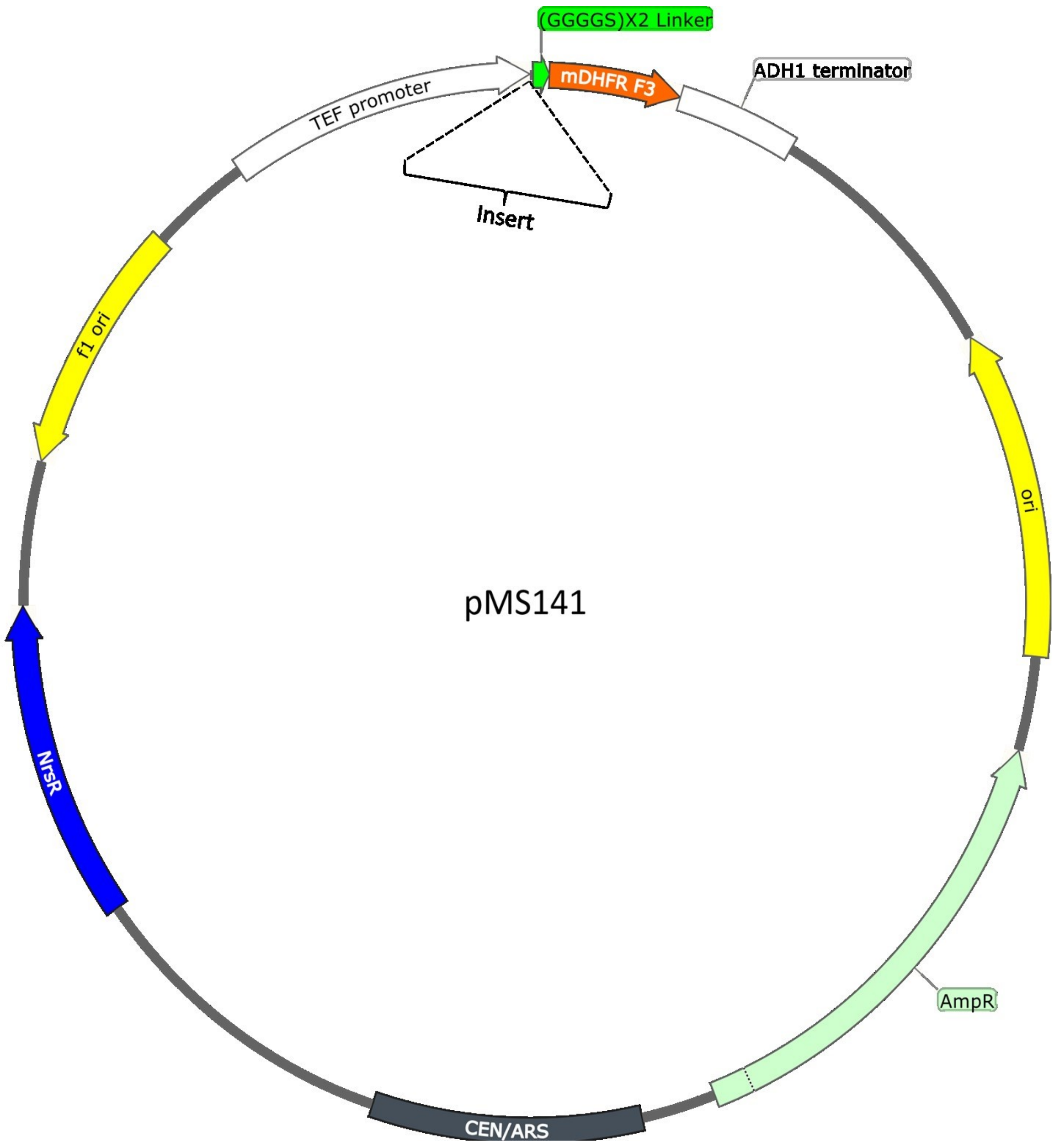


**Figure S2. Maps of the constructs used in the split mDHFR PCA.** Human C1 subunit insertion sites in each plasmid are indicated.

Supplementary Figure S3

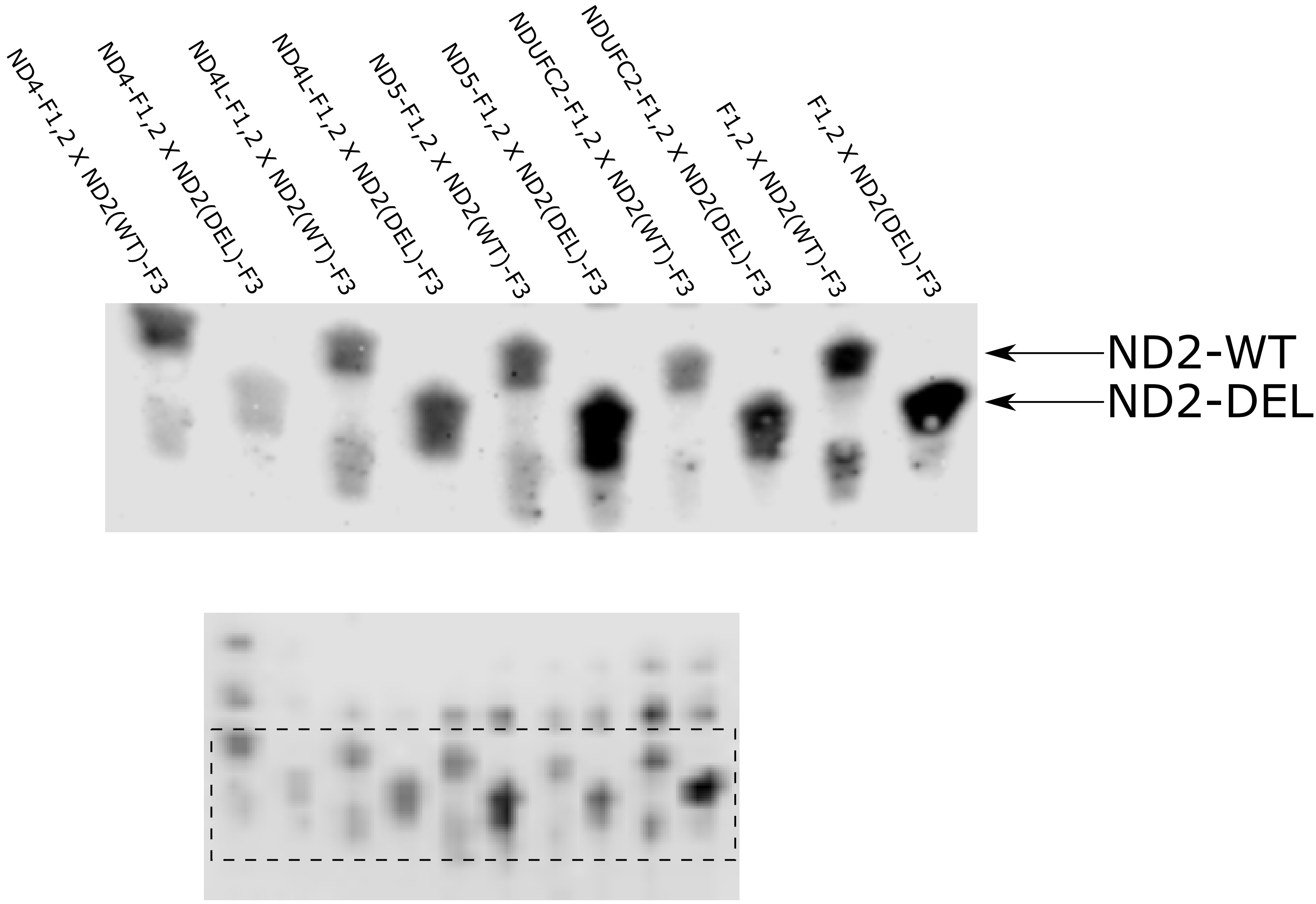


**Figure S3. Validation of protein expression of the ND2 frameshift mutant and the wild type protein.** Western blot representing three independent mating experiments of either wild type ND2 or the ND2 frameshift mutant with all four tested interactors. Bands representing the wild type and the frameshift mutant are indicated. Complementary full western blot is shown at the lower panel. Dashed rectangle represent the area from which the upper panel was cropped.

**Supplementary Table S1 – C1 subunits in the two split mDHFR libraries.** +/- indicates successful/unsuccessful cloning, respectively.

| Subunit | p340 library (mDHFR F1,2) | pMS-141 library (mDHFR F3) |
|---|---|---|
| ND1 | + | + |
| ND2 | + | + |
| ND3 | + | + |
| ND4 | + | + |
| ND4L | + | + |
| ND5 | + | + |
| ND6 | - | + |
| NDUFA1 | + | + |
| NDUFA2 | + | + |
| NDUFA3 | + | + |
| NDUFA5 | + | + |
| NDUFA6 | + | + |
| NDUFA7 | + | + |
| NDUFA8 | + | + |
| NDUFA9 | - | + |
| NDUFA10 | + | + |
| NDUFA11 | + | + |
| NDUFA12 | - | + |
| NDUFA13 | + | + |
| NDUFAB1 | - | + |
| NDUFB1 | + | + |
| NDUFB2 | + | + |
| NDUFB3 | + | + |
| NDUFB4 | + | + |
| NDUFB5 | + | + |
| NDUFB6 | + | + |
| NDUFB7 | + | + |
| NDUFB8 | + | + |
| NDUFB9 | - | + |
| NDUFB10 | + | + |
| NDUFB11 | + | + |
| NDUFC1 | + | + |
| NDUFC2 | + | + |
| NDUFS1 | - | + |
| NDUFS2 | + | + |
| NDUFS3 | + | + |
| NDUFS4 | - | + |
| NDUFS5 | + | + |
| NDUFS6 | + | + |
| NDUFS7 | + | + |
| NDUFS8 | + | + |
| NDUFV1 | + | + |
| NDUFV2 | + | + |
| NDUFV3 | + | + |

**Supplementary Table S2 – Description of the yeast media used in this work**

| Medium | Description |
|---|---|
| SD agar plates | 2.2% w/v Bacto-agar, 2% w/v Dextrose, 0.67% w/v Bacto-yeast nitrogen base without amino acids and with ammonium sulfate (Difco) and with vital amino and nucleic acids as described in (Sherman, 2002). |
| SD-His agar plates | 2.2% w/v Bacto-agar, 2% w/v Dextrose, 0.67% w/v Bacto-yeast nitrogen base without amino acids and with ammonium sulfate (Difco) and with vital amino and nucleic acids as described in (Sherman, 2002), excluding Histidine. |
| SD-His agar plates + cloNAT | 2.2% w/v Bacto-agar, 2% w/v Dextrose, 0.67% w/v Bacto-yeast nitrogen base without amino acids, supplemented with ammonium sulfate, vital amino and nucleic acids, excluding Histidine, as previously described (Sherman, 2002) and 0.2mg\ml cloNAT. |
| SD-His liquid medium + cloNAT | 2% w/v Dextrose, 0.67% w/v Bacto-yeast nitrogen base w/o amino acids and with ammonium sulfate (Difco), with added vital amino and nucleic acids, excluding Histidine, as previously described (Sherman, 2002) and supplemented with 0.2mg\ml cloNAT. |
| SD agar plates + MTX | 4% w/v Bacto-agar (NOBLE), 2% w/v Dextrose 0.67% w/v Bacto-yeast nitrogen base w/o amino acids and with ammonium sulfate (Difco), supplemented with vital amino and nucleic acids as previously described (Sherman, 2002) and 0.2% w/v MTX. |
| SD liquid medium + MTX | 2% w/v Dextrose 0.67% w/v Bacto-yeast nitrogen base w/o amino acids and with ammonium sulfate (Difco), supplemented with vital amino and nucleic acids as previously described (Sherman, 2002) and 0.2% w/v MTX. |
| YPD agar plates | 2% Bacto-agar, 2% Dextrose, 2% Bacto-Peptone, 1% Bacto-yeast extract, tittered by HCl, to pH=5.8-6.0. |
| YPD agar plates + cloNAT | 2% Bacto-agar, 2% Dextrose, 2% Bacto-Peptone, 1% Bacto-yeast extract, supplemented by 0.2mg\ml cloNAT and tittered by HCl, to pH=5.8-6.0. |

**Supplementary Table S3 – Complex I source of PCR template used for insert preparation**

| Subunit | source |
|---|---|
| NDUFA1 | Skeletal Muscle (Ambion cat no. 7982) |
| NDUFA2 | Brain (Ambion cat no. 7962) |
| NDUFA3 | Brain (Ambion cat no. 7962) |
| NDUFA5 | Brain (Ambion cat no. 7962) |
| NDUFA6 | Brain (Ambion cat no. 7962) |
| NDUFA7 | Brain (Ambion cat no. 7962) |
| NDUFA8 | Brain (Ambion cat no. 7962) |
| NDUFA9 | Brain (Ambion cat no. 7962) |
| NDUFA10 | Pancreas (Ambion cat no. 7954) |
| NDUFA11 | Brain (Ambion cat no. 7962) |
| NDUFA12 | Brain (Ambion cat no. 7962) |
| NDUFA13 | Brain (Ambion cat no. 7962) |
| NDUFAB1 | Brain (Ambion cat no. 7962) |
| NDUFB1 | Brain (Ambion cat no. 7962) |
| NDUFB2 | Brain (Ambion cat no. 7962) |
| NDUFB3 | Brain (Ambion cat no. 7962) |
| NDUFB4 | Brain (Ambion cat no. 7962) |
| NDUFB5 | Pancreas (Ambion cat no. 7954) |
| NDUFB6 | Brain (Ambion cat no. 7962) |
| NDUFB7 | Brain (Ambion cat no. 7962) |
| NDUFB8 | Pancreas (Ambion cat no. 7954) |
| NDUFB9 | Pancreas (Ambion cat no. 7954) |
| NDUFB10 | Brain (Ambion cat no. 7962) |
| NDUFB11 | Heart (Ambion cat no. 7966) |
| NDUFC1 | Pancreas (Ambion cat no. 7954) |
| NDUFC2 | Skeletal Muscle (Ambion cat no. 7982) |
| NDUFS1 | Brain (Ambion cat no. 7962) |
| NDUFS2 | Heart (Ambion cat no. 7966) |
| NDUFS3 | Pancreas (Ambion cat no. 7954) |
| NDUFS4 | Heart (Ambion cat no. 7966) |
| NDUFS5 | Heart (Ambion cat no. 7966) |
| NDUFS6 | Brain (Ambion cat no. 7962) |
| NDUFS7 | Pancreas (Ambion cat no. 7954) |
| NDUFS8 | Pancreas (Ambion cat no. 7954) |
| NDUFV1 | Pancreas (Ambion cat no. 7954) |
| NDUFV2 | Pancreas (Ambion cat no. 7954) |
| NDUFV3 | Brain (Ambion cat no. 7962) |

**Supplementary Table S4– The cytoplasmic translation conversion of human mtDNA encoded complex I subunits.**

| Subunit | Converted coding sequence (W/O stop codon) |
|---|---|
| ND1 | ATGCCAATGGCAAATCTGCTGCTCCTCATCGTGCCAATCCTGATCGCCATGGCCTTCCTCATGCT<br>GACTGAAAGAAAAATTCTGGGATACATGCAGCTCAGGAAGGGGCCTAACGTGGTGGGACCTTA<br>TGGACTGCTCCAGCCCTTTGCTGATGCTATGAAGCTGTTCACAAAAGAGCCCCTGAAACCAGCC<br>ACCTCTACAATCACCCTGTACATTACCGCTCCTACCCTGGCTCTGACAATTGCCCTGCTGCTGTGG<br>ACCCCTCTCCCTATGCCAAATCCTCTGGTGAACCTGAATCTGGGCCTCCTCTTTATCCTGGCCACC<br>AGCAGCCTGGCCGTGTACTCCATCCTGTGGAGCGGATGGGCTTCTAACAGCAATTACGCCCTGA<br>TCGGTGCCCTGAGGGCCGTGGCCCAGACCATTTCTTACGAGGTGACCCTCGCCATTATCCTGCTC<br>TCAACCCTGCTGATGAGCGGCTCTTTCAACCTCTCAACCCTGATTACAACCCAGGAGCACCTCTG<br>GCTGCTCCTCCCCAGCTGGCCACTGGCCATGATGTGGTTTATCAGCACCCTGGCTGAGACAAAC<br>CGGACCCCCTTTGATCTGGCTGAGGGCGAGTCTGAGCTGGTCTCCGGATTCAATATTGAGTACG<br>CAGCAGGGCCATTCGCTCTGTTCTTCATGGCCGAGTATACAAATATTATTATGATGAACACACTG<br>ACTACTACTATCTTCCTGGGTACTACATACGATGCTCTGAGTCCCGAACTCTACACCACTTACTTC<br>GTGACCAAAACCCTGCTGCTGACTAGCCTGTTCCTGTGGATCAGGACCGCCTATCCACGATTCCG<br>ATACGACCAGCTGATGCATCTGCTGTGGAAGAACTTCCTGCCACTCACCCTGGCTCTGCTCATGT<br>GGTACGTGAGTATGCCAATCACTATCAGCTCTATCCCTCCACAGACC |
| ND2 | ATGAATCCCCTGGCCCAACCCGTCATCTACTCTACCATCTTTGCAGGCACACTCATCACA<br>GCGCTAAGCTCGCACTGGTTTTTTACCTGGGTAGGCCTAGAAATGAACATGCTAGCTTT<br>TATTCCAGTTCTAACCAAAAAAATGAACCCTCGTTCCACAGAAGCTGCCATCAAGTATT<br>TCCTCACGCAAGCAACCGCATCCATGATCCTTCTAATGGCTATCCTCTTCAACAATATGC<br>TCTCCGGACAATGGACCATGACCAATACTACCAATCAATACTCATCATTAATGATCATG<br>ATGGCTATGGCAATGAAACTAGGAATGGCCCCCTTTCACTTCTGGGTCCCAGAGGTTA<br>CCCAAGGCACCCCTCTGACATCCGGCCTGCTTCTTCTCACATGGCAAAAACTAGCCCCC<br>ATCTCAATCATGTACCAAATCTCTCCCTCACTAAACGTAAGCCTTCTCCTCACTCTCTCA<br>ATCTTATCCATCATGGCAGGCAGTTGGGGTGGATTAAACCAAACCCAGCTACGCAAAA<br>TCTTAGCATACTCCTCAATTACCCACATGGGATGGATGATGGCAGTTCTACCGTACAAC<br>CCTAACATGACCATTCTTAATTTAACTATTTATATTATCCTAACTACTACCGCATTCCTAC<br>TACTCAACTTAAACTCCAGCACCACGACCCTACTACTATCTCGCACCTGGAACAAGCTA<br>ACATGGCTAACACCCTTAATTCCATCCACCCTCCTCTCCCTAGGAGGCCTGCCCCCGCTA<br>ACCGGCTTTTTGCCCAAATGGGCCATTATCGAAGAATTCACAAAAAACAATAGCCTCAT<br>CATCCCCACCATCATGGCCACCATCACCCTCCTTAACCTCTACTTCTACCTACGCCTAAT<br>CTACTCCACCTCAATCACACTACTCCCCATGTCTAACAACGTAAAAATGAAATGGCAGT<br>TTGAACATACAAAACCCACCCCATTCCTCCCCACACTCATCGCCCTTACCACGCTACTCC<br>TACCTATCTCCCCTTTTATGCTAATGATCTTA |
| ND2-DEL | ATGAATCCCCTGGCCCAACCCGTCATCTACTCTACCATCTTTGCAGGCACACTCATCACA<br>GCGCTAAGCTCGCACTGGTTTTTTACCTGGGTAGGCCTAGAAATGAACATGCTAGCTTT<br>TATTCCAGTTCTAACCAAAAAAATGAACCCTCGTTCCACAGAAGCTGCCATCAAGTATT<br>TCCTCACGCAAGCAACCGCATCCATGATCCTTCTAATGGCTATCCTCTTCAACAATATGC<br>TCTCCGGACAATGGACCATGACCAATACTACCAATCAATACTCATCATTAATGATCATG<br>ATGGCTATGGCAATGAAACTAGGAATGGCCCCCTTTCACTTCTGGGTCCCAGAGGTTA<br>CCCAAGGCACCCCTCTGACATCCGGCCTGCTTCTTCTCACATGGCAAAAACTAGCCCCC |

| | |
|---|---|
| | ATCTCAATCATGTACCAAATCTCTCCCTCACTAAACGTAAGCCTTCTCCTCACTCTCTCA ATCTTATCCATCATGGCAGGCAGTTGGGGTGGATTAAACCAAACCCAGCTACGCAAAA TCTTAGCATACTCCTCAATTACCCACATGGGATGGATGATGGCAGTTCTACCGTACAAC CCTAACATGACCATTCTTAATTTAACTATTTATATTATCCTAACTACTACCGCATTCCTAC TACTCAACCTGTTGCAACACCACGACCCGACGACTATTAGTCATTTGAAGCAGGCCAAC ATGACTAATACTCTTAACTCAATACACCCCCCATTACCA |
| ND3 | ATGAACTTCGCCTTAATCTTAATGATCAACACCCTCCTAGCCTTACTACTAATGATCATC ACATTTTGGCTACCACAACTCAACGGCTACATGGAAAAATCCACCCCTTACGAGTGCG GCTTCGACCCTATGTCCCCCGCCCGCGTCCCTTTCTCCATGAAATTCTTCTTAGTAGCTA TCACCTTCTTATTATTTGATCTAGAAATCGCCCTCCTTTTACCCCTACCATGGGCCCTACA AACAACTAACCTGCCACTAATGGTTATGTCATCCCTCTTATTAATCATCATCCTAGCCCT AAGTCTGGCCTATGAGTGGCTACAAAAAGGATTAGACTGGGCCGAA |
| ND4 | ATGCTAAAACTAATCGTCCCAACAATCATGTTACTACCACTGACATGGCTTTCCAAAAA ACACATGATCTGGATCAACACAACCACCCACAGCCTAATCATCAGCATCATCCCTCTAC TATTTTTTAACCAAATCAACAACAACCTATTTAGCTGTTCCCCAACCTTTTCCTCCGACCC CCTAACAACCCCCCTCCTAATGCTAACTACCTGGCTCCTACCCCTCACAATCATGGCAA GCCAACGCCACTTATCCAGTGAACCACTATCACGAAAAAAACTCTACCTCTCTATGCTA ATCTCCCTACAAATCTCCTTAATCATGACATTCACAGCCACAGAACTAATCATGTTTTAT ATCTTCTTCGAAACCACACTTATCCCCACCTTGGCTATCATCACCCGATGGGGCAACCA GCCAGAACGCCTGAACGCAGGCACATACTTCCTATTCTACACCCTAGTAGGCTCCCTTC CCCTACTCATCGCACTAATCTACACTCACAACACCCTAGGCTCACTAAACATCCTACTAC TCACTCTCACTGCCCAAGAACTATCAAACTCCTGGGCCAACAACTTAATGTGGCTAGCT TACACAATGGCTTTTATGGTAAAGATGCCTCTTTACGGACTCCACTTATGGCTCCCTAA AGCCCATGTCGAAGCCCCCATCGCTGGGTCAATGGTACTTGCCGCAGTACTCTTAAAA CTAGGCGGCTATGGTATGATGCGCCTCACACTCATCCTCAACCCCCTGACAAAACACAT GGCCTACCCCTTCCTTGTACTATCCCTATGGGGCATGATCATGACAAGCTCCATCTGCC TACGACAAACAGACCTAAAATCGCTCATCGCATACTCTTCAATCAGCCACATGGCCCTC GTAGTAACAGCCATCCTCATCCAAACCCCCTGGAGCTTCACCGGCGCAGTCATCCTCAT GATCGCCCACGGGCTTACATCCTCATTACTATTCTGCCTAGCAAACTCAAACTACGAAC GCACTCACAGTCGCATCATGATCCTCTCTCAAGGACTTCAAACTCTACTCCCACTAATG GCTTTTTGGTGGCTTCTAGCAAGCCTCGCTAACCTCGCCTTACCCCCCACTATCAACCTA CTGGGAGAACTCTCTGTGCTAGTAACCACGTTCTCCTGGTCAAATATCACTCTCCTACTT ACAGGACTCAACATGCTAGTCACAGCCCTATACTCCCTCTACATGTTTACCACAACACA ATGGGGCTCACTCACCCACCACATCAACAACATGAAACCCTCATTCACACGAGAAAAC ACCCTCATGTTCATGCACCTATCCCCCATCCTCCTCCTATCCCTCAACCCCGACATCATCA CCGGGTTTTCCTCT |
| ND4L | ATGCCCCTCATTTACATGAATATTATGCTAGCATTTACCATCTCACTTCTAGGAATGCTA GTATATCGCTCACACCTCATGTCCTCCCTACTATGCCTAGAAGGAATGATGCTATCGCT GTTCATTATGGCTACTCTCATGACCCTCAACACCCACTCCCTCTTAGCCAATATTGTGCC TATTGCCATGCTAGTCTTTGCCGCCTGCGAAGCAGCGGTGGGCCTAGCCCTACTAGTCT CAATCTCCAACACATATGGCCTAGACTACGTACATAACCTAAACCTACTCCAATGC |

| | |
|---|---|
| ND5 | ATGACCATGCACACTACTATGACCACCCTAACCCTGACTTCCCTAATTCCCCCATCCTT<br>ACCACCCTCGTTAACCCTAACAAAAAAACTCATACCCCCATTATGTAAAATCCATTGTC<br>GCATCCACCTTTATTATCAGTCTCTTCCCCACAACAATGTTCATGTGCCTAGACCAAGAA<br>GTTATTATCTCGAACTGGCACTGGGCCACAACCCAAACAACCCAGCTCTCCCTAAGCTT<br>CAAACTAGACTACTTCTCCATGATGTTCATCCCTGTAGCATTGTTCGTTACATGGTCCAT<br>CATGGAATTCTCACTGTGGTATATGAACTCAGACCCAAACATTAATCAGTTCTTCAAAT<br>ATCTACTCATTTTCCTAATTACCATGCTAATCTTAGTTACCGCTAACAACCTATTCCAACT<br>GTTCATCGGCTGGGAGGGCGTAGGAATTATGTCCTTCTTGCTCATCAGTTGGTGGTAC<br>GCCCGAGCAGATGCCAACACAGCAGCCATTCAAGCAGTCCTATACAACCGTATCGGCG<br>ATATCGGTTTCATCCTCGCCTTAGCATGGTTTATCCTACACTCCAACTCATGGGACCCAC<br>AACAAATGGCCCTTCTAAACGCTAATCCAAGCCTCACCCCACTACTAGGCCTCCTCCTA<br>GCAGCAGCAGGCAAATCAGCCCAATTAGGTCTCCACCCCTGGCTCCCCTCAGCCATGG<br>AAGGCCCCACCCCAGTCTCAGCCCTACTCCACTCAAGCACTATGGTTGTAGCAGGAATC<br>TTCTTACTCATCCGCTTCCACCCCCTAGCAGAAAATAGCCCACTAATCCAAACTCTAACA<br>CTATGCTTAGGCGCTATCACCACTCTGTTCGCAGCAGTCTGCGCCCTTACACAAAATGA<br>CATCAAAAAAATCGTAGCCTTCTCCACTTCAAGTCAACTAGGACTCATGATGGTTACAA<br>TCGGCATCAACCAACCACACCTAGCATTCCTGCACATCTGTACCCACGCCTTCTTCAAA<br>GCCATGCTATTTATGTGCTCCGGGTCCATCATCCACAACCTTAACAATGAACAAGATAT<br>TCGAAAAATGGGAGGACTACTCAAAACCATGCCTCTCACTTCAACCTCCCTCACCATTG<br>GCAGCCTAGCATTAGCAGGAATGCCTTTCCTCACAGGTTTCTACTCCAAAGACCACATC<br>ATCGAAACCGCAAACATGTCATACACAAACGCCTGGGCCCTATCTATTACTCTCATCGC<br>TACCTCCCTGACAAGCGCCTATAGCACTCGAATGATTCTTCTCACCCTAACAGGTCAAC<br>CTCGCTTCCCCACCCTTACTAACATTAACGAAAATAACCCCACCCTACTAAACCCCATTA<br>AACGCCTGGCAGCCGGAAGCCTATTCGCAGGATTTCTCATTACTAACAACATTTCCCCC<br>GCATCCCCCTTCCAAACAACAATCCCCCTCTACCTAAAACTCACAGCCCTCGCTGTCACT<br>TTCCTAGGACTTCTAACAGCCCTAGACCTCAACTACCTAACCAACAAACTTAAAATGAA<br>ATCCCCACTATGCACATTTTATTTCTCCAACATGCTCGGATTCTACCCTAGCATCACACA<br>CCGCACAATCCCCTATCTAGGCCTTCTTACGAGCCAAAACCTGCCCCTACTCCTCCTAGA<br>CCTAACCTGGCTAGAAAAGCTATTACCTAAAACAATTTCACAGCACCAAATCTCCACCT<br>CCATCATCACCTCAACCCAAAAAGGCATGATTAAACTTTACTTCCTCTCTTTCTTCTTCCC<br>ACTCATCCTAACCCTACTCCTAATCACATAA |
| ND6 | ATGATGTATGCATTATTTTTATTATCTGTTGGTTTAGTTATGGGTTTTGTAGGTTTTTCAT<br>CAAAGCCTAGTCCAATATATGGTGGTTTGGTTTTGATTGTTAGTGGTGTTGTCGGTTGT<br>GTTATTATCTTAAACTTCGGTGGTGGTTACATGGGTTTGATGGTTTTCTTGATATACTTG<br>GGTGGTATGATGGTTGTTTTTGGTTATACCACAGCTATGGCCATTGAAGAATATCCAG<br>AAGCATGGGGTTCTGGTGTCGAAGTTTTAGTAAGTGTTTTGGTTGGTTTGGCAATGGA<br>AGTTGGTTTGGTTTTATGGGTTAAAGAATACGATGGTGTCGTTGTCGTTGTTAACTTCA<br>ATAGTGTCGGTTCTTGGATGATCTACGAAGGTGAAGGTTCTGGTTTAATTAGAGAAGA<br>TCCTATTGGTGCTGGTGCCTTATACGATTACGGTAGATGGTTAGTCGTAGTTACAGGTT<br>GGACCTTATTCGTTGGTGTTTATATTGTTATTGAAATCGCTAGAGGTAAT |

**Supplementary Table S5 - Primers used for amplifications of the inserts of C1 subunits with 5' and 3' tails for cloning into p340 and pMS141 plasmids**

| Primer name | Primer Sequence |
|---|---|
| ND1 into pMS141 forward | GGAGTACTTGTTTTTAGAATATACGGTCAACGAACTATAATTAACTAAACATGCCAATGGCAAATCTGCTGCTCCTCATC |
| ND1 into p340 forward | TCTTTTTTTTAGTTTTAAAACACCAGAACTTAGTTTCGACGGATTCTAGAATGCCAATGGCAAATCTGCTGCTCCTCATC |
| ND1 into p340 reverse | TTCTTCACCTTTAGACATAGAGCCACCTCCACCAGATCCGCCACCGCCGGTCTGTGGAGGGATAGAGCTGATAGTGATTG |
| ND1 into pMS141 reverse | CATGTCTACTTTACTCATAGAGCCACCTCCACCAGATCCGCCACCGCCGGTCTGTGGAGGGATAGAGCTGATAGTGATTG |
| ND2 into p340 forward | TCTTTTTTTTAGTTTTAAAACACCAGAACTTAGTTTCGACGGATTCTAGAATGAATCCCCTGGCCCAACCCGTCATCTAC |
| ND2 into p340 reverse | AATTCTTCACCTTTAGACATAGAGCCACCTCCACCAGATCCGCCACCGCCTAAGATCATTAGCATAAAAGGGGAGATAGG |
| ND2 into pMS141 forward | GGAGTACTTGTTTTTAGAATATACGGTCAACGAACTATAATTAACTAAACATGAATCCCCTGGCCCAACCCGTCATCTAC |
| ND2 into pMS141 reverse | ACCATGTCTACTTTACTCATAGAGCCACCTCCACCAGATCCGCCACCGCCTAAGATCATTAGCATAAAAGGGGAGATAGG |
| ND3 into p340 forward | TCTTTTTTTTAGTTTTAAAACACCAGAACTTAGTTTCGACGGATTCTAGAATGAACTTCGCCTTAATCTTAATGATCAAC |
| ND3 into p340 reverse | ATTCTTCACCTTTAGACATAGAGCCACCTCCACCAGATCCGCCACCGCCTTCGGCCCAGTCTAATCCTTTTTGTAGCCAC |
| ND3 into pMS141 forward | GGAGTACTTGTTTTTAGAATATACGGTCAACGAACTATAATTAACTAAACATGAACTTCGCCTTAATCTTAATGATCAAC |
| ND3 into pMS141 reverse | CCATGTCTACTTTACTCATAGAGCCACCTCCACCAGATCCGCCACCGCCTTCGGCCCAGTCTAATCCTTTTTGTAGCCAC |
| ND4 into p340 forward | TCTTTTTTTTAGTTTTAAAACACCAGAACTTAGTTTCGACGGATTCTAGAATGCTAAAACTAATCGTCCCAACAATCATG |
| ND4 into p340 reverse | ATTCTTCACCTTTAGACATAGAGCCACCTCCACCAGATCCGCCACCGCCAGAGGAAAACCCGGTGATGATGTCGGGGTTG |
| ND4 into pMS141 forward | GGAGTACTTGTTTTTAGAATATACGGTCAACGAACTATAATTAACTAAACATGCTAAAACTAATCGTCCCAACAATCATG |
| ND4 into pMS141 reverse | CCATGTCTACTTTACTCATAGAGCCACCTCCACCAGATCCGCCACCGCCAGAGGAAAACCCGGTGATGATGTCGGGGTTG |
| ND4L into p340 forward | CTTTTTTTTAGTTTTAAAACACCAGAACTTAGTTTCGACGGATTCTAGAATGCCCCTCATCTACATGAATATCATGCTAG |
| ND4L into p340 reverse | ATTCTTCACCTTTAGACATAGAGCCACCTCCACCAGATCCGCCACCGCCGCATTGGAGTAGGTTTAGGTTATGTACGTAG |
| ND4L into pMS141 forward | GAGTACTTGTTTTTAGAATATACGGTCAACGAACTATAATTAACTAAACATGCCCCTCATCTACATGAATATCATGCTAG |
| ND4L into pMS141 reverse | CCATGTCTACTTTACTCATAGAGCCACCTCCACCAGATCCGCCACCGCCGCATTGGAGTAGGTTTAGGTTATGTACGTAG |
| ND5 into p340 forward | TTTTTTTTAGTTTTAAAACACCAGAACTTAGTTTCGACGGATTCTAGAATGACCATGCACACTACTATGACCACCCTAAC |
| ND5 into p340 reverse | AATTCTTCACCTTTAGACATAGAGCCACCTCCACCAGATCCGCCACCGCCTGTGATTAGGAGTAGGGTTAGGATGAGTGG |
| ND5 into pMS141 forward | AGTACTTGTTTTTAGAATATACGGTCAACGAACTATAATTAACTAAACATGACCATGCACACTACTATGACCACCCTAAC |
| ND5 into pMS141 reverse | ACCATGTCTACTTTACTCATAGAGCCACCTCCACCAGATCCGCCACCGCCTGTGATTAGGAGTAGGGTTAGGATGAGTGG |
| ND6 into p340 forward | CTTTTTTTTAGTTTTAAAACACCAGAACTTAGTTTCGACGGATTCTAGAATGATGTATGCATTATTTTTATTATCTGTTG |
| ND6 into p340 reverse | TCTTCACCTTTAGACATAGAGCCACCTCCACCAGATCCGCCACCGCCATTACCTCTAGCGATTTCAATAACAATATAAAC |

| | |
|---|---|
| ND6 into pMS141 forward | GAGTACTTGTTTTTAGAATATACGGTCAACGAACTATAATTAACTAAACATGATGTATGCATTATTTTTATTATCTGTTG |
| ND6 into pMS141 reverse | ATGTCTACTTTACTCATAGAGCCACCTCCACCAGATCCGCCACCGCCATTACCTCTAGCGATTTCAATAACAATATAAAC |
| NDUFA1 into p340 forward | TCTTTTTTTTAGTTTTAAAACACCAGAACTTAGTTTCGACGGATTCTAGAATGTGGTTCGAGATTCTCCCCGGACTCTC |
| NDUFA1 into p340 reverse | ATTCTTCACCTTTAGACATGAGCCACCTCCACCAGATCCGCCACCGCCATCAATGTTCTCCAAACCCTTTGACACATAG |
| NDUFA1 into pMS141 forward | GGAGTACTTGTTTTTAGAATATACGGTCAACGAACTATAATTAACTAAACATGTGGTTCGAGATTCTCCCCGGACTCTC |
| NDUFA1 into pMS141 reverse | CCATGTCTACTTTACTCATAGAGCCACCTCCACCAGATCCGCCACCGCCATCAATGTTCTCCAAACCCTTTGACACATAG |
| NDUFA2 into p340 forward | TCTTTTTTTTAGTTTTAAAACACCAGAACTTAGTTTCGACGGATTCTAGAATGGCGGCGGCCGCAGCAAGTCG |
| NDUFA2 into p340 reverse | AATTCTTCACCTTTAGACATAGAGCCACCTCCACCAGATCCGCCACCGCCGGCTTTACCACTTAGAACGTTCTCCAG |
| NDUFA2 into pMS141 forward | GGAGTACTTGTTTTTAGAATATACGGTCAACGAACTATAATTAACTAAACATGGCGGCGGCCGCAGCAAGTCG |
| NDUFA2 into pMS141 reverse | ACCATGTCTACTTTACTCATAGAGCCACCTCCACCAGATCCGCCACCGCCGGCTTTACCACTTAGAACGTTCTCCAG |
| NDUFA3 into p340 forward | TCTTTTTTTTAGTTTTAAAACACCAGAACTTAGTTTCGACGGATTCTAGAATGGCTGCGAGAGTCGGCGCCTTCCTCAAG |
| NDUFA3 into p340 reverse | AATTCTTCACCTTTAGACATAGAGCCACCTCCACCAGATCCGCCACCGCCCAGTTTCTTCAGCCACTCCAGGCTGGG |
| NDUFA3 into pMS141 forward | GGAGTACTTGTTTTTAGAATATACGGTCAACGAACTATAATTAACTAAACATGGCTGCGAGAGTCGGCGCCTTCCTCAAG |
| NDUFA3 into pMS141 reverse | ACCATGTCTACTTTACTCATAGAGCCACCTCCACCAGATCCGCCACCGCCCAGTTTCTTCAGCCACTCCAGGCTGGG |
| NDUFA5 into p340 forward | TCTTTTTTTTAGTTTTAAAACACCAGAACTTAGTTTCGACGGATTCTAGAATGGCGGGTGTGCTGAAGAAGACCACTGGC |
| NDUFA5 into p340 reverse | AATTCTTCACCTTTAGACATAGAGCCACCTCCACCAGATCCGCCACCGCCTATTGGCCATTTCCACTGATCGGCAGGAGG |
| NDUFA5 into pMS141 forward | GGAGTACTTGTTTTTAGAATATACGGTCAACGAACTATAATTAACTAAACATGGCGGGTGTGCTGAAGAAGACCACTGGC |
| NDUFA5 into pMS141 reverse | ACCATGTCTACTTTACTCATAGAGCCACCTCCACCAGATCCGCCACCGCCTATTGGCCATTTCCACTGATCGGCAGGAGG |
| NDUFA6 into p340 forward | TCTTTTTTTTAGTTTTAAAACACCAGAACTTAGTTTCGACGGATTCTAGAATGGGAAAAGACATTCGCCCG |
| NDUFA6 into p340 reverse | AATTCTTCACCTTTAGACATAGAGCCACCTCCACCAGATCCGCCACCGCCTGGATCGTGGCCAACATAGAAC |
| NDUFA6 into pMS141 forward | GGAGTACTTGTTTTTAGAATATACGGTCAACGAACTATAATTAACTAAACATGGGAAAAGACATTCGCCCG |
| NDUFA6 into pMS141 reverse | ACCATGTCTACTTTACTCATAGAGCCACCTCCACCAGATCCGCCACCGCCTGGATCGTGGCCAACATAGAAC |
| NDUFA7 into p340 forward | TCTTTTTTTTAGTTTTAAAACACCAGAACTTAGTTTCGACGGATTCTAGAATGGCGTCCGCCACCCGTCTC |
| NDUFA7 into p340 reverse | AATTCTTCACCTTTAGACATAGAGCCACCTCCACCAGATCCGCCACCGCCCAGGTAAGGCTGGTCCGAGGACAGCTC |
| NDUFA7 into pMS141 forward | GGAGTACTTGTTTTTAGAATATACGGTCAACGAACTATAATTAACTAAACATGGCGTCCGCCACCCGTCTC |
| NDUFA7 into pMS141 reverse | ACCATGTCTACTTTACTCATAGAGCCACCTCCACCAGATCCGCCACCGCCCAGGTAAGGCTGGTCCGAGGACAGCTC |
| NDUFA8 into p340 forward | CTTTTTTTTAGTTTTAAAACACCAGAACTTAGTTTCGACGGATTCTAGAATGCCGGGGATAGTGGAGCTGCCCACTCTAG |
| NDUFA8 into p340 reverse | AATTCTTCACCTTTAGACATAGAGCCACCTCCACCAGATCCGCCACCGCCCTTGGTCCAGAAATAAAAGCGGCTGCCATG |

| | |
|---|---|
| NDUFA8 into pMS141 forward | GAGTACTTGTTTTTAGAATATACGGTCAACGAACTATAATTAACTAAACATGCCGGGGATAGTGGAGCTGCCCACTCTAG |
| NDUFA8 into pMS141 reverse | ACCATGTCTACTTTACTCATAGAGCCACCTCCACCAGATCCGCCACCGCCCTTGGTCCAGAAATAAAAGCGGCTGCCATG |
| NDUFA9 into p340 forward | TCTTTTTTTTAGTTTTAAAACACCAGAACTTAGTTTCGACGGATTCTAGACTTGGTCCAGAAATAAAAGCGGCTGCCATG |
| NDUFA9 into p340 reverse | AATTCTTCACCTTTAGACATAGAGCCACCTCCACCAGATCCGCCACCGCCAATGTTGACGGTCTTGGCCGGCTTCACATC |
| NDUFA9 into pMS141 forward | GGAGTACTTGTTTTTAGAATATACGGTCAACGAACTATAATTAACTAAACATGGCGGCTGCCGCACAATCCCGGGTTGTC |
| NDUFA9 into pMS141 reverse | ACCATGTCTACTTTACTCATAGAGCCACCTCCACCAGATCCGCCACCGCCAATGTTGACGGTCTTGGCCGGCTTCACATC |
| NDUFA10 into p340 forward | TCTTTTTTTTAGTTTTAAAACACCAGAACTTAGTTTCGACGGATTCTAGAATGGCCTTGCGGCTCCTGAAG |
| NDUFA10 into p340 reverse | AATTCTTCACCTTTAGACATAGAGCCACCTCCACCAGATCCGCCACCGCCCTTCAGCCAGATCCACTTGTCTCCC |
| NDUFA10 into pMS141 forward | GGAGTACTTGTTTTTAGAATATACGGTCAACGAACTATAATTAACTAAACATGGCCTTGCGGCTCCTGAAG |
| NDUFA10 into pMS141 reverse | ACCATGTCTACTTTACTCATAGAGCCACCTCCACCAGATCCGCCACCGCCCTTCAGCCAGATCCACTTGTCTCCC |
| NDUFA11 into p340 forward | TCTTTTTTTTAGTTTTAAAACACCAGAACTTAGTTTCGACGGATTCTAGAATGGCGCCGAAGGTTTTTCGTCAGTAC |
| NDUFA11 into p340 reverse | AATTCTTCACCTTTAGACATAGAGCCACCTCCACCAGATCCGCCACCGCCCACCTTGGGTTTTGCAAACACCTCCC |
| NDUFA11 into pMS141 forward | GGAGTACTTGTTTTTAGAATATACGGTCAACGAACTATAATTAACTAAACATGGCGCCGAAGGTTTTTCGTCAGTAC |
| NDUFA11 into pMS141 reverse | ACCATGTCTACTTTACTCATAGAGCCACCTCCACCAGATCCGCCACCGCCCACCTTGGGTTTTGCAAACACCTCCC |
| NDUFA12 into p340 forward | TCTTTTTTTTAGTTTTAAAACACCAGAACTTAGTTTCGACGGATTCTAGAATGGAGTTAGTGCAGGTCCTGAAAC |
| NDUFA12 into p340 reverse | AATTCTTCACCTTTAGACATAGAGCCACCTCCACCAGATCCGCCACCGCCCTTGTAAGGTGTTGAAGGTGGGATC |
| NDUFA12 into pMS141 forward | GGAGTACTTGTTTTTAGAATATACGGTCAACGAACTATAATTAACTAAACATGGAGTTAGTGCAGGTCCTGAAAC |
| NDUFA12 into pMS141 reverse | ACCATGTCTACTTTACTCATAGAGCCACCTCCACCAGATCCGCCACCGCCCTTGTAAGGTGTTGAAGGTGGGATC |
| NDUFA13 into p340 forward | TCTTTTTTTTAGTTTTAAAACACCAGAACTTAGTTTCGACGGATTCTAGAATGGCGGCGTCAAAGGTGAAGCAG |
| NDUFA13 into p340 reverse | AATTCTTCACCTTTAGACATAGAGCCACCTCCACCAGATCCGCCACCGCCCGTGTACCACATGAAGCCGTGGCTG |
| NDUFA13 into pMS141 forward | GGAGTACTTGTTTTTAGAATATACGGTCAACGAACTATAATTAACTAAACATGGCGGCGTCAAAGGTGAAGCAG |
| NDUFA13 into pMS141 reverse | ACCATGTCTACTTTACTCATAGAGCCACCTCCACCAGATCCGCCACCGCCCGTGTACCACATGAAGCCGTGGCTG |
| NDUFAB1 into P340 forward | TCTTTTTTTTAGTTTTAAAACACCAGAACTTAGTTTCGACGGATTCTAGAATGGCGTCTCGTGTCCTTTCAG |
| NDUFAB1 into p340 reverse | AATTCTTCACCTTTAGACATAGAGCCACCTCCACCAGATCCGCCACCGCCTTCATATACATCCTTCTTATCTGCAATG |
| NDUFAB1 into pMS141 reverse | ACCATGTCTACTTTACTCATAGAGCCACCTCCACCAGATCCGCCACCGCCTTCATATACATCCTTCTTATCTGCAATG |
| NDUFAB1into pMS141 forward | GGAGTACTTGTTTTTAGAATATACGGTCAACGAACTATAATTAACTAAACATGGCGTCTCGTGTCCTTTCAG |
| NDUFB1 into p340 forward | TCTTTTTTTTAGTTTTAAAACACCAGAACTTAGTTTCGACGGATTCTAGAATGATTTGCTGGCGTCACCC |
| NDUFB1 into p340 reverse | AATTCTTCACCTTTAGACATAGAGCCACCTCCACCAGATCCGCCACCGCCCTTCCAGGTAACTTCTTCACTGGG |

| | |
|---|---|
| NDUFB1 into pMS141 forward | GGAGTACTTGTTTTTAGAATATACGGTCAACGAACTATAATTAACTAAACATGATTTGCTGGCGTCACCC |
| NDUFB1 into pMS141 reverse | ACCATGTCTACTTTACTCATAGAGCCACCTCCACCAGATCCGCCACCGCCCTTCCAGGTAACTTCTTCACTGGG |
| NDUFB2 into p340 forward | TCTTTTTTTTAGTTTTAAAACACCAGAACTTAGTTTCGACGGATTCTAGAATGTCCGCTCTGACTCGGCTG |
| NDUFB2 into p340 reverse | AATTCTTCACCTTTAGACATAGAGCCACCTCCACCAGATCCGCCACCGCCGTCTTCATCATCAGGAGGGATACCTAATTC |
| NDUFB2 into pMS141 forward | GGAGTACTTGTTTTTAGAATATACGGTCAACGAACTATAATTAACTAAACATGTCCGCTCTGACTCGGCTG |
| NDUFB2 into pMS141 reverse | ACCATGTCTACTTTACTCATAGAGCCACCTCCACCAGATCCGCCACCGCCGTCTTCATCATCAGGAGGGATACCTAATTC |
| NDUFB3 into p340 forward | TCTTTTTTTTAGTTTTAAAACACCAGAACTTAGTTTCGACGGATTCTAGAATGGCCCATGAACATGGACATG |
| NDUFB3 into p340 reverse | AATTCTTCACCTTTAGACATAGAGCCACCTCCACCAGATCCGCCACCGCCGTGATGCTTCTTATCTTTATTCAGGGACTC |
| NDUFB3 into pMS141 forward | GGAGTACTTGTTTTTAGAATATACGGTCAACGAACTATAATTAACTAAACATGGCCCATGAACATGGACATG |
| NDUFB3 into pMS141 reverse | ACCATGTCTACTTTACTCATAGAGCCACCTCCACCAGATCCGCCACCGCCGTGATGCTTCTTATCTTTATTCAGGGACTC |
| NDUFB4 into p340 forward | TCTTTTTTTTAGTTTTAAAACACCAGAACTTAGTTTCGACGGATTCTAGAATGTCGTTCCCAAAGTATAAGCCG |
| NDUFB4 into p340 reverse | AATTCTTCACCTTTAGACATAGAGCCACCTCCACCAGATCCGCCACCGCCATATGAGAGGTGAAATGTTCGATCC |
| NDUFB4 into pMS141 forward | GGAGTACTTGTTTTTAGAATATACGGTCAACGAACTATAATTAACTAAACATGTCGTTCCCAAAGTATAAGCCG |
| NDUFB4 into pMS141 reverse | ACCATGTCTACTTTACTCATAGAGCCACCTCCACCAGATCCGCCACCGCCATATGAGAGGTGAAATGTTCGATCC |
| NDUFB5 into p340 forward | TCTTTTTTTTAGTTTTAAAACACCAGAACTTAGTTTCGACGGATTCTAGAATGGCGGCCATGAGTTTGTTG |
| NDUFB5 into p340 reverse | AATTCTTCACCTTTAGACATAGAGCCACCTCCACCAGATCCGCCACCGCCATTGTCAGGAGTTGCTTTCGGAGAATG |
| NDUFB5 into pMS141 forward | GGAGTACTTGTTTTTAGAATATACGGTCAACGAACTATAATTAACTAAACATGGCGGCCATGAGTTTGTTG |
| NDUFB5 into pMS141 reverse | ACCATGTCTACTTTACTCATAGAGCCACCTCCACCAGATCCGCCACCGCCATTGTCAGGAGTTGCTTTCGGAGAATG |
| NDUFB6 into p340 forward | TCTTTTTTTTAGTTTTAAAACACCAGAACTTAGTTTCGACGGATTCTAGAATGACGGGGTACACTCCGGATG |
| NDUFB6 into p340 reverse | AATTCTTCACCTTTAGACATAGAGCCACCTCCACCAGATCCGCCACCGCCATGATGTTGATCAGGAAATTCTTTCATTGG |
| NDUFB6 into pMS141 forward | GGAGTACTTGTTTTTAGAATATACGGTCAACGAACTATAATTAACTAAACATGACGGGGTACACTCCGGATG |
| NDUFB6 into pMS141 reverse | ACCATGTCTACTTTACTCATAGAGCCACCTCCACCAGATCCGCCACCGCCATGATGTTGATCAGGAAATTCTTTCATTGG |
| NDUFB7 into p340 forward | TCTTTTTTTTAGTTTTAAAACACCAGAACTTAGTTTCGACGGATTCTAGAATGGGGGCCCACCTGGTCCG |
| NDUFB7 into p340 reverse | AATTCTTCACCTTTAGACATAGAGCCACCTCCACCAGATCCGCCACCGCCCAGGGCCACCTTGGGGTCCACTTC |
| NDUFB7 into pMS141 forward | GGAGTACTTGTTTTTAGAATATACGGTCAACGAACTATAATTAACTAAACATGGGGGCCCACCTGGTCCG |
| NDUFB7 into pMS141 reverse | ACCATGTCTACTTTACTCATAGAGCCACCTCCACCAGATCCGCCACCGCCCAGGGCCACCTTGGGGTCCACTTC |
| NDUFB8 into p340 forward | TCTTTTTTTTAGTTTTAAAACACCAGAACTTAGTTTCGACGGATTCTAGAATGGCGGTGGCCAGGGCCGG |
| NDUFB8 into p340 reverse | AATTCTTCACCTTTAGACATAGAGCCACCTCCACCAGATCCGCCACCGCCGATCTCATAGTGAACCACCCGCTCTGGTTC |

| | |
|---|---|
| NDUFB8 into pMS141 forward | GGAGTACTTGTTTTTAGAATATACGGTCAACGAACTATAATTAACTAAACATGGCGGTGGCCAGGGCCGG |
| NDUFB8 into pMS141 reverse | ACCATGTCTACTTTACTCATAGAGCCACCTCCACCAGATCCGCCACCGCCGATCTCATAGTGAACCACCCGCTCTGGTTC |
| NDUFB9 into p340 forward | TCTTTTTTTTAGTTTTAAAACACCAGAACTTAGTTTCGACGGATTCTAGAATGGCGTTCTTGGCGTCGGG |
| NDUFB9 into p340 reverse | AATTCTTCACCTTTAGACATAGAGCCACCTCCACCAGATCCGCCACCGCCCATGGGCCGCTCCCGGGGTC |
| NDUFB9 into pMS141 forward | GGAGTACTTGTTTTTAGAATATACGGTCAACGAACTATAATTAACTAAACATGGCGTTCTTGGCGTCGGG |
| NDUFB9 into pMS141 reverse | ACCATGTCTACTTTACTCATAGAGCCACCTCCACCAGATCCGCCACCGCCCATGGGCCGCTCCCGGGGTC |
| NDUFB10 into p340 forward | TCTTTTTTTTAGTTTTAAAACACCAGAACTTAGTTTCGACGGATTCTAGAATGCCGGACAGCTGGGACAAGGATGTGTAC |
| NDUFB10 into p340 reverse | AATTCTTCACCTTTAGACATAGAGCCACCTCCACCAGATCCGCCACCGCCGGAGGTGGCAGCGGCGGCCTCTTTTGCAGC |
| NDUFB10 into pMS141 forward | GGAGTACTTGTTTTTAGAATATACGGTCAACGAACTATAATTAACTAAACATGCCGGACAGCTGGGACAAGGATGTGTAC |
| NDUFB10 into pMS141 reverse | ACCATGTCTACTTTACTCATAGAGCCACCTCCACCAGATCCGCCACCGCCGGAGGTGGCAGCGGCGGCCTCTTTTGCAGC |
| NDUFB11 into p340 forward | TCTTTTTTTTAGTTTTAAAACACCAGAACTTAGTTTCGACGGATTCTAGAATGGCGGCTGGGCTGTTTGG |
| NDUFB11 into p340 reverse | AATTCTTCACCTTTAGACATAGAGCCACCTCCACCAGATCCGCCACCGCCCTCATCCTCTGGCAGCTGGATCTTG |
| NDUFB11 into pMS141 forward | GGAGTACTTGTTTTTAGAATATACGGTCAACGAACTATAATTAACTAAACATGGCGGCTGGGCTGTTTGG |
| NDUFB11 into pMS141 reverse | ACCATGTCTACTTTACTCATAGAGCCACCTCCACCAGATCCGCCACCGCCCTCATCCTCTGGCAGCTGGATCTTG |
| NDUFC1 into p340 forward | TCTTTTTTTTAGTTTTAAAACACCAGAACTTAGTTTCGACGGATTCTAGAATGGCGCCGTCCGCCTTGCTG |
| NDUFC1 into p340 reverse | AATTCTTCACCTTTAGACATAGAGCCACCTCCACCAGATCCGCCACCGCCTTCCAGCCCATTTCTTCTTTTGTACTC |
| NDUFC1 into pMS141 forward | GGAGTACTTGTTTTTAGAATATACGGTCAACGAACTATAATTAACTAAACATGGCGCCGTCCGCCTTGCTG |
| NDUFC1 into pMS141 reverse | ACCATGTCTACTTTACTCATAGAGCCACCTCCACCAGATCCGCCACCGCCTTCCAGCCCATTTCTTCTTTTGTACTC |
| NDUFC2 into pMS141 forward | GAGTACTTGTTTTTAGAATATACGGTCAACGAACTATAATTAACTAAACATGATCGCACGGCGGAACCCAGAACCCTTAC |
| NDUFC2 into pMS141 reverse | CCATGTCTACTTTACTCATAGAGCCACCTCCACCAGATCCGCCACCGCCACGTATTGGATGGAATTTTTCAAAAATTTC |
| NDUFC2 into p340 forward | CTTTTTTTTAGTTTTAAAACACCAGAACTTAGTTTCGACGGATTCTAGAATGATCGCACGGCGGAACCCAGAACCCTTAC |
| NDUFC2 into p340 reverse | AATTCTTCACCTTTAGACATGAGCCACCTCCACCAGATCCGCCACCGCCACGTATTGGATGGAATTTTTCAAAAATTTC |
| NDUFS1 into p340 forward | TCTTTTTTTTAGTTTTAAAACACCAGAACTTAGTTTCGACGGATTCTAGAATGTTAAGGATACCTGTAAGAAAGGCC |
| NDUFS1 into p340 reverse | AATTCTTCACCTTTAGACATAGAGCCACCTCCACCAGATCCGCCACCGCCGCATATGGATGGTTCCTCTACTGC |
| NDUFS1 into pMS141 forward | GGAGTACTTGTTTTTAGAATATACGGTCAACGAACTATAATTAACTAAACATGTTAAGGATACCTGTAAGAAAGGCC |
| NDUFS1 into pMS141 reverse | ACCATGTCTACTTTACTCATAGAGCCACCTCCACCAGATCCGCCACCGCCGCATATGGATGGTTCCTCTACTGC |
| NDUFS2 into p340 forward | TCTTTTTTTTAGTTTTAAAACACCAGAACTTAGTTTCGACGGATTCTAGAATGGCGGCGCTGAGGGCTTTG |
| NDUFS2 into p340 reverse | AATTCTTCACCTTTAGACATAGAGCCACCTCCACCAGATCCGCCACCGCCCCGATCTACTTCTCCAAATACAATATCTTG |

| | |
|---|---|
| NDUFS2 into pMS141 forward | GGAGTACTTGTTTTTAGAATATACGGTCAACGAACTATAATTAACTAAACATGGCGGCGCTGAGGGCTTTG |
| NDUFS2 into pMS141 reverse | ACCATGTCTACTTTACTCATAGAGCCACCTCCACCAGATCCGCCACCGCCCCGATCTACTTCTCCAAATACAATATCTTG |
| NDUFS3 into p340 forward | TCTTTTTTTTAGTTTTAAAACACCAGAACTTAGTTTCGACGGATTCTAGAATGGCGGCGGCGGCGGTAGC |
| NDUFS3 into p340 reverse | AATTCTTCACCTTTAGACATAGAGCCACCTCCACCAGATCCGCCACCGCCCTTGGCATCAGGCTTCTTGTCTCCGGCTTC |
| NDUFS3 into pMS141 forward | GGAGTACTTGTTTTTAGAATATACGGTCAACGAACTATAATTAACTAAACATGGCGGCGGCGGCGGTAGC |
| NDUFS3 into pMS141 reverse | ACCATGTCTACTTTACTCATAGAGCCACCTCCACCAGATCCGCCACCGCCCTTGGCATCAGGCTTCTTGTCTCCGGCTTC |
| NDUFS4 into p340 forward | TCTTTTTTTTAGTTTTAAAACACCAGAACTTAGTTTCGACGGATTCTAGAATGGCGGCGGTGTCAATGTC |
| NDUFS4 into p340 reverse | AATTCTTCACCTTTAGACATAGAGCCACCTCCACCAGATCCGCCACCGCCTTTTGTGGATACTCTTGTTCTTTTGTTCC |
| NDUFS4 into pMS141 forward | GGAGTACTTGTTTTTAGAATATACGGTCAACGAACTATAATTAACTAAACATGGCGGCGGTGTCAATGTC |
| NDUFS4 into pMS141 reverse | ACCATGTCTACTTTACTCATAGAGCCACCTCCACCAGATCCGCCACCGCCTTTTGTGGATACTCTTGTTCTTTTGTTCC |
| NDUFS5 into p340 forward | TCTTTTTTTTAGTTTTAAAACACCAGAACTTAGTTTCGACGGATTCTAGAATGCCTTTCTTGGACATCCAGAAAAGGTTC |
| NDUFS5 into p340 reverse | AATTCTTCACCTTTAGACATAGAGCCACCTCCACCAGATCCGCCACCGCCGGGCCGAGGCTCCCCCTTGC |
| NDUFS5 into pMS141 forward | GGAGTACTTGTTTTTAGAATATACGGTCAACGAACTATAATTAACTAAACATGCCTTTCTTGGACATCCAGAAAAGGTTC |
| NDUFS5 into pMS141 reverse | ACCATGTCTACTTTACTCATAGAGCCACCTCCACCAGATCCGCCACCGCCGGGCCGAGGCTCCCCCTTGC |
| NDUFS6 into p340 forward | TCTTTTTTTTAGTTTTAAAACACCAGAACTTAGTTTCGACGGATTCTAGAATGGCGGCGGCGATGACCTTCTGCCGGCTG |
| NDUFS6 into p340 reverse | ATTCTTCACCTTTAGACATAGAGCCACCTCCACCAGATCCGCCACCGCCGTGGTGGTGCTGTCTGAACTGGAGCCCACAG |
| NDUFS6 into pMS141 forward | GGAGTACTTGTTTTTAGAATATACGGTCAACGAACTATAATTAACTAAACATGGCGGCGGCGATGACCTTCTGCCGGCTG |
| NDUFS6 into pMS141 reverse | CCATGTCTACTTTACTCATAGAGCCACCTCCACCAGATCCGCCACCGCCGTGGTGGTGCTGTCTGAACTGGAGCCCACAG |
| NDUFS7 into p340 forward | TCTTTTTTTTAGTTTTAAAACACCAGAACTTAGTTTCGACGGATTCTAGAATGGCGGTGCTGTCAGCTCC |
| NDUFS7 into p340 reverse | AATTCTTCACCTTTAGACATAGAGCCACCTCCACCAGATCCGCCACCGCCCCTGCGGTACCAGATCTGCAGC |
| NDUFS7 into pMS141 forward | GGAGTACTTGTTTTTAGAATATACGGTCAACGAACTATAATTAACTAAACATGGCGGTGCTGTCAGCTCC |
| NDUFS7 into pMS141 reverse | ACCATGTCTACTTTACTCATAGAGCCACCTCCACCAGATCCGCCACCGCCCCTGCGGTACCAGATCTGCAGC |
| NDUFS8 into p340 forward | TCTTTTTTTTAGTTTTAAAACACCAGAACTTAGTTTCGACGGATTCTAGAATGCGCTGCCTGACCACGCC |
| NDUFS8 into p340 reverse | AATTCTTCACCTTTAGACATAGAGCCACCTCCACCAGATCCGCCACCGCCCCGATACAAGTAGTCAGCCTGGATGTTGGC |
| NDUFS8 into pMS141 forward | GGAGTACTTGTTTTTAGAATATACGGTCAACGAACTATAATTAACTAAACATGCGCTGCCTGACCACGCC |
| NDUFS8 into pMS141 reverse | ACCATGTCTACTTTACTCATAGAGCCACCTCCACCAGATCCGCCACCGCCCCGATACAAGTAGTCAGCCTGGATGTTGGC |
| NDUFV1 into p340 forward | TCTTTTTTTTAGTTTTAAAACACCAGAACTTAGTTTCGACGGATTCTAGAATGCTGGCAACACGGCGGCTG |
| NDUFV1 into p340 reverse | AATTCTTCACCTTTAGACATAGAGCCACCTCCACCAGATCCGCCACCGCCAGAGGCAGCCTGCCGGGCCTG |

| | |
|---|---|
| NDUFV1 into pMS141 forward | GGAGTACTTGTTTTTAGAATATACGGTCAACGAACTATAATTAACTAAACATGCTGGCAACACGGCGGCTG |
| NDUFV1 into pMS141 reverse | ACCATGTCTACTTTACTCATAGAGCCACCTCCACCAGATCCGCCACCGCCAGAGGCAGCCTGCCGGGCCTG |
| NDUFV2 into p340 forward | TCTTTTTTTTAGTTTTAAAACACCAGAACTTAGTTTCGACGGATTCTAGAATGTTCTTCTCCGCGGCGCTC |
| NDUFV2 into p340 reverse | AATTCTTCACCTTTAGACATAGAGCCACCTCCACCAGATCCGCCACCGCCAAGGCCTGCTTGTACACCAAATCCAG |
| NDUFV2 into pMS141 forward | GGAGTACTTGTTTTTAGAATATACGGTCAACGAACTATAATTAACTAAACATGTTCTTCTCCGCGGCGCTC |
| NDUFV2 into pMS141 reverse | ACCATGTCTACTTTACTCATAGAGCCACCTCCACCAGATCCGCCACCGCCAAGGCCTGCTTGTACACCAAATCCAG |
| NDUFV3 into p340 forward | CTTTTTTTTAGTTTTAAAACACCAGAACTTAGTTTCGACGGATTCTAGAATGGCTGCCCCGTGTTTGCTGCGGCAAGGAC |
| NDUFV3 into p340 reverse | AATTCTTCACCTTTAGACATAGAGCCACCTCCACCAGATCCGCCACCGCCGTGTCGAGGTGACTCCCGGCCTGAGGAGGG |
| NDUFV3 into pMS141 forward | GAGTACTTGTTTTTAGAATATACGGTCAACGAACTATAATTAACTAAACATGGCTGCCCCGTGTTTGCTGCGGCAAGGAC |
| NDUFV3 into pMS141 reverse | ACCATGTCTACTTTACTCATAGAGCCACCTCCACCAGATCCGCCACCGCCGTGTCGAGGTGACTCCCGGCCTGAGGAGGG |

**Supplementary Table S6 - Primers used for validation of successful insertion of C1 subunits to p340 and pMS141**

| Primer Name | Primer Sequence |
|---|---|
| pMS141 ins val forward | GGAGTACTTGTTTTTAGAATATACGGTCAACG |
| pMS-41 ins val reverse | AAGTCAGGCGATCACACATTAAAAGCTATAC |
| p340 ins val forward | TGACTAATAAGTATATAAAGACGGTAGGTATTG |
| p340 ins val reverse | TAGTACAAATCAATTTTAAGGTCAATTTACC |

**Supplementary Table S7 – C1 subunits RefSeq protein accession numbers**

| Subunit | RefSeq Accession Number |
|---|---|
| ND1 | YP_003024026.1 |
| ND2 | YP_003024027.1 |
| ND3 | YP_003024033.1 |
| ND4 | YP_003024035.1 |
| ND4L | YP_003024034.1 |
| ND5 | YP_003024036.1 |
| ND6 | YP_003024037.1 |
| NDUFA1 | NP_004532.1 |
| NDUFA2 | NP_002479.1 |
| NDUFA3 | NP_004533.1 |
| NDUFA5 | NP_004991.1 |
| NDUFA6 | NP_002481.2 |
| NDUFA7 | NP_004992.2 |
| NDUFA8 | NP_055037.1 |
| NDUFA9 | NP_004993.1 |
| NDUFA10 | NP_004535.1 |
| NDUFA11 | NP_783313.1 |
| NDUFA12 | NP_061326.1 |
| NDUFA13 | NP_057049.5 |
| NDUFAB1 | NP_004994.1 |
| NDUFB1 | NP_004536.2 |
| NDUFB2 | NP_004537.1 |
| NDUFB3 | NP_002482.1 |
| NDUFB4 | NP_004538.2 |
| NDUFB5 | NP_002483.1 |
| NDUFB6 | NP_002484.1 |
| NDUFB7 | NP_004137.2 |
| NDUFB8 | NP_004995.1 |
| NDUFB9 | NP_004996.1 |
| NDUFB10 | NP_004539.1 |
| NDUFB11 | NP_061929.2 |
| NDUFC1 | NP_002485.1 |
| NDUFC2 | NP_004540.1 |
| NDUFS1 | NP_004997.4 |
| NDUFS2 | NP_004541.1 |
| NDUFS3 | NP_004542.1 |
| NDUFS4 | NP_002486.1 |
| NDUFS5 | NP_004543.1 |
| NDUFS6 | NP_002484.1 |
| NDUFS7 | NP_077718.3 |
| NDUFS8 | NP_002487.1 |
| NDUFV1 | NP_009034.2 |
| NDUFV2 | NP_066552.2 |
| NDUFV3 | NP_001001503.1 |
